# Discovery of Natural MCL1 Inhibitors using Pharmacophore modelling, QSAR, Docking, ADMET, Molecular Dynamics, and DFT Analysis

**DOI:** 10.1101/2024.10.14.618373

**Authors:** Uddalak Das, Tathagata Chanda, Jitendra Kumar, Anitha Peter

## Abstract

Mcl-1, a Bcl-2 family protein, is a key regulator of apoptosis and is often overexpressed in cancers such as lung, breast, pancreatic, cervical, ovarian cancers, leukemia, and lymphoma. Its role in inhibiting apoptosis enables tumor cells to evade cell death and contributes to drug resistance. Targeting Mcl-1 is crucial in inducing apoptosis and overcoming resistance to therapies that target other anti-apoptotic proteins, making it a prominent target for anticancer drug development across multiple malignancies. So, our study aimed to discover potent antileukemic compounds targeting MCL1. We began by selecting a diverse set of molecules from the BindingDB database to construct a structure-based pharmacophore model, which was subsequently used to virtually screen a library of 407,270 compounds from the COCONUT database. Subsequently, we developed an e-pharmacophore model using the co-crystallized inhibitor AMG-176, for further screening. A QSAR model was then implemented to estimate the IC_50_ values of these compounds, filtering those with predicted IC_50_ values below the median. The top hits were subjected to molecular docking and MMGBSA binding energy calculations against MCL1, leading to the selection of two promising candidates for further ADMET analysis. To assess their electronic properties, density functional theory (DFT) calculations were conducted, which included geometry optimization, frontier molecular orbital (FMO) analysis, HOMO-LUMO gaps, and global reactivity descriptors. These analyses confirmed favourable profiles and reactivity for the chosen compounds. In addition, predictions for physicochemical and ADMET properties aligned well with the expected bioactivity and safety profiles of the candidates. Molecular dynamics (MD) simulations further validated their strong binding affinity and stability, positioning them as potential MCL1 inhibitors. Our comprehensive computational approach highlights these compounds as promising and safe antileukemic agents, with future *in vivo* and *in vitro* validation recommended for further confirmation.

## INTRODUTION

Cancer remains a pervasive global health challenge, responsible for millions of deaths annually across a spectrum of over 200 distinct diseases (Thun *et al*., 2010). To comprehend and combat this complexity, cancer biologists have delineated several organizing principles known as cancer hallmarks (Hanahan & Weinberg, 2011). Among these, resistance to cell death, particularly apoptosis, stands out as a critical feature acquired by neoplastic cells during tumorigenesis due to genomic instability and dysregulation of cell cycle checkpoints (Ferguson *et al*., 2015).

Apoptosis evasion is a hallmark of cancer progression, enabling malignant cells to evade natural cellular mechanisms aimed at eliminating abnormal or damaged cells (Fernald & Kurokawa, 2013). Dysregulation of apoptotic signalling pathways, mediated in part by the Bcl-2 family of proteins, leads to sustained cell proliferation and tumor development (Qian *et al*., 2022). Among these proteins, Mcl-1 (Myeloid cell leukemia 1) emerges as a pivotal regulator implicated in the resistance of cancer cells to chemotherapy and radiotherapy (Sancho *et al*., 2022).

The Bcl-2 family comprises both anti-apoptotic (e.g., Bcl-2, Bcl-xL, Mcl-1) and pro-apoptotic (e.g., Bak, Bax, BH3-only proteins) members, which intricately regulate the apoptotic process (Hardwick & Soane, 2013). Mcl-1, known for its high expression in various cancer tissues compared to other anti-apoptotic Bcl-2 family members, plays a crucial role in preventing apoptosis by inhibiting mitochondrial outer membrane permeabilization (MOMP) and cytochrome C release (Morciano *et al*., 2016). Its upregulation not only promotes cancer cell survival but also confers resistance to conventional anti-cancer therapies (Sulkshane & Teni, 2022).

Efforts to counteract Mcl-1’s anti-apoptotic effects have identified it as a promising therapeutic target in cancer treatment (Tantawy *et al*., 2023). Despite progress in identifying Mcl-1 inhibitors with potent anti-cancer effects *in vitro*, challenges such as poor *in vivo* pharmacokinetic properties hinder their clinical translation (Tantawy *et al*., 2023).

Computer-aided drug design (CADD) has emerged as a crucial tool in overcoming these challenges, offering efficient screening and optimization of potential Mcl-1 inhibitors (Baig *et al*., 2018; Gurung *et al*., 2021; Iwaloye *et al*., 2023; Macalino *et al*., 2015; Niazi & Mariam, 2023). By integrating pharmacophore modeling, molecular docking, molecular dynamics simulations, and ADME prediction, researchers can rationally design novel compounds that selectively target Mcl-1, enhancing efficacy and minimizing off-target effects.

In recent years, natural products have garnered attention as a valuable source of lead compounds for drug discovery. Their structural diversity and bioactivity make them attractive candidates for the development of novel therapeutics (Newman & Cragg, 2020). Integration of natural products with computational screening techniques presents an opportunity to identify potent FLT3 inhibitors from large compound libraries, such as the Collection of Open Natural Products (COCONUT) database (Sorokina *et al*., 2021).

In this study, we aimed to design novel Mcl-1 inhibitors using computational methods and fragments from the Enamine database. Our approach encompasses pharmacophore modeling, fragment linking, molecular docking, binding free energy calculations, ADME prediction, and molecular dynamics simulations to identify potential candidates with improved pharmacological profiles.

## 2. MATERIALS AND METHODS

### 2.1. Dataset generation and screening

The COCONUT (version 2022, Collection of Open Natural ProdUcTs Online) database was utilized as the screening source, containing approximately 4,07,270 molecules. The molecules were then filtered using QikProp module in “Structure Filtering” option Schrödinger 2023-1, with zero violation of Lipinski’s Rule of 5 generating an output of 2,76,409 molecules. The filtered structures where then prepared using the LigPrep Module with the OPLS-2005 force field. The possible states of molecules were generated using Epik at pH 7.0 ± 2.0, retained specified chirality, followed by tautomer generation and the generation of up to 16 low-energy conformations per ligand (Johnston *et al*., 2023). The total number of generated structures were around ∼3.57 million.

### 2.2. Pharmacophore modelling/screening

To ensure a comprehensive analysis of ligand activity and structural diversity, two distinct pharmacophore models were developed and utilized for screening. Initially, a diverse ligands-based pharmacophore model was created using ligands, allowing for the capture of varied ligand activities across different structural types. This model ensured a broad exploration of potential ligand interactions. Following this, an e-Pharmacophore model was generated, to enhance specificity. The combination of these two pharmacophore models enabled a dual screening approach that leveraged the diverse ligand activities and structures from the diverse ligands-based model while refining the search with the highly specific e-Pharmacophore model. This strategy maximized the identification of potential ligands by balancing broad coverage with detailed specificity (Giordano *et al*., 2022).

#### 2.2.1. Diverse-ligand based pharmacophore

A pharmacophore comprises distinct characteristics like steric properties, hydrogen bond donor (HBD), hydrogen bond acceptor (HBA), and electronic chemical attributes. These features indicate how a compound functions uniquely within the active biological site. (Islam *et. al.,* 2024) The development of ligand-based pharmacophore design relied on identified actives with confirmed pharmacological effects targeting the chosen receptor.

To generate the pharmacophore model, an initial subset comprising the first 100 compounds linked to the MCL1 was extracted from the binding database, sorted based on their respective IC_50_ values. (Ganji *et. al.,* 2023) This curated dataset was the foundation for subsequent pharmacophore model construction and screening endeavors. Within the LigPrep module, the OPLS4 force field was engaged (Lu *et al.,* 2021), concomitant with default parameters for ionization. Furthermore, routine procedures encompassing desalination, tautomerization, and computational adjustments were implemented per software defaults. It helps to prepare high-quality 3D structures for drug-like molecules.

In the Develop Pharmacophore Model module, the hypothesis match was set to 75% and the number of features in the hypothesis was kept from 4 to 7 with the preferred number to 5. The ranking and scoring of the hypothesis were set to the default “Phase Hypo Score” (Yu *et. al.,* 2021). The Generate Conformer and Minimize Output conformer options were activated, with the target number of conformers set to 50 (Cole *et. al.,* 2018).

The prepared ligands were screened using the best generated pharmacophore model, using the “Phase Module”. Prefer partial matches of more features were activated. All the pharmacophore properties were selected for screening.

##### 2.2.2. E-pharmacophore

The e-pharmacophore model of MCL1 was constructed using the PDB ID 6O6F (Chain A), where the co-crystallized AMG-176 inhibitor was docked. Subsequently, the e-pharmacophore model was developed utilizing PHASE v.3 (Schrödinger Release, 2023-1). The model parameters included a maximum of 7 features, with donors defined as vectors. The minimum feature-feature distance was set to 2 Å, while the minimum distance for features of the same group type was 4 Å. A receptor-based excluded volume shell was activated, with radii set to the Van der Waals radii of receptor atoms and a fixed scaling factor of 0.5. The excluded volume shell thickness was limited to 5 Å. The generated e-pharmacophore was used to screen the compounds furthur.

##### 2.2.3. Validation of the pharmacophore models

The developed pharmacophore model was verified to check its ability to predict the activity of new compounds effectively. This procedure is a prerequisite before the pharmacophore model can be employed for virtual screening (Kaserer *et al*., 2015). The parameters used for evaluating the efficiency of the developed pharmacophore model are enrichment factor (EF), receiver operating characteristic (ROC) curves (Triballeau *et al*., 2005), Boltzmann-enhanced discrimination of ROC (BEDROC) (Truchon & Bayly, 2007), and robust initial enhancement (RIE) (Truchon & Bayly, 2007).

A decoy set was created using the Generate DUDE Decoys program, which is found at http://dude.docking.org/generate (Mysinger *et al*., 2012). For converting the output into 3D structures, Open Babel v2.4.1 was used (O’Boyle *et al*., 2011). Ligand preparation was done following the default settings and the protocols of LigPrep. The OPLS4 force field has been employed in the minimization procedure (Lu *et al*., 2021).

### 2.3. QSAR Analysis

AutoQSAR module of Schrödinger was used to build the QSAR model and predict the IC_50_ of the screened compounds. The model was built with the top 100 MCL1 inhibitors from the BindingDB database (Liu *et al*., 2007), sorted by IC_50_ values. Ligand preparation resulted in 367 compounds. The dataset was divided into 75% training (192 compounds) and 25% test sets (64 compounds) keeping 111 compounds (30%) as validation set. Model parameters included a maximum correlation of 0.80 between independent variables along with activation of binary fingerprints, and numerical descriptors. The best model was used to predict the IC_50_ of the 2,024 screened compounds and screen compounds below the median IC_50_ value.

### 2.4. Molecular Interactions

#### 2.4.1. Protein preparation

The X-ray crystallographic structure of the MCL1 target protein [PDB ID: 6O6F], found at https://www.rcsb.org, with a resolution of 1.60 Å was used in the study (Caenepeel *et al*., 2018). The structure analysis was carried out using the PDBsum web server (Laskowski *et al*., 2018) (https://www.ebi.ac.uk/thornton-srv/databases/pdbsum). PDBsum online server was also used to check the validation of the MCL1 with the Ramachandran plot (Ramachandran *et al*., 1963).

In the Schrödinger Maestro protein preparation wizard, the protein was pre-processed with the PROPKA module for an optimization of H-bonds (Olsson *et al*., 2011), followed by minimization of structures towards convergence of heavy atoms at RMSD 0.3Å using OPLS4 force field and removal of water molecules more than 5Å away from ligands afterward (Mahdizadeh *et al*., 2021).

#### 2.4.2. Receptor grid generation

The receptor grid was generated keeping the hydrophobic region and also the region where the inhibitor AMG-176 was attached to the complex, at the centroid of the grid. The coordinates of the receptor grid were X=-19.36, Y=-7.55, Z=-19.24, with ligand size upto 18Å

#### 2.4.3. Molecular docking

Docking was limited to ligands with 100 rotatable bonds and fewer than 500 atoms. Van der Waals radii scaling factor was set to 0.80, with a partial charge cutoff of 0.15. Sample nitrogen inversions and sample ring conformations were activated, and the ligand sampling was set to flexible. All predicted functional groups had bias sampling of torsions enabled. The module was configured to promote intramolecular hydrogen bonds and improve conjugated pi groups’ planarity. Extra precision (XP) docking was done using the Glide module of Schrödinger. The 2D and 3D visualization of the protein-ligand interations were done using PyMOL v3.0.5 and BIOVIA Discovery Studio Visualizer v24.1.0.23298

#### 2.4.4. MM-GBSA Screening

MM-GBSA (Molecular Mechanics Generalized Born Surface Area) analysis demonstrates a stronger correlation with real-life wet lab experiments compared to traditional docking methods in *in silico* drug discovery (Taylor & Ho, 2023). This is primarily due to its ability to account for the dynamic nature of molecular interactions, including the contributions of solvation effects, entropic factors, and the inherent flexibility of both ligands and target proteins.

The top 10% of the ligands screened through XP docking were analysed using MM-GBSA. The prime module was utilized to predict the energy parameters obtained from the MM-GBSA (Molecular Mechanics Generalized Born Surface Area) simulation. It aimed to predict the amounts of the stabilization energy coming from the potential interaction between the selected ligands and the target receptor 6O0K. The VGSB solvation model was used and the force field was set as OPLS4 (Jawarkar *et al*., 2022).

### 2.5. Density Functional Theory (DFT) Analysis

Gaussian 09 and GaussView molecular visualization software were employed to perform theoretical Density Functional Theory (DFT) calculations, aimed at studying the electronic properties and chemical reactivity of the selected compounds. The geometry of the compounds was optimized using the B3LYP functional (Tirado-Rives & Jorgensen, 2008) in conjunction with the 6-31G (Rassolov *et al*., 2001) basis set. Key quantum descriptors were then calculated to provide a comprehensive analysis of the compounds’ behaviour. These descriptors include the highest occupied molecular orbital (HOMO), lowest unoccupied molecular orbital (LUMO), and the HOMO-LUMO energy gap (ΔE = E_LUMO - E_HOMO). The dipole moment (μ) and total energy (E_total) were also determined, reflecting the overall stability and polarity of the molecules.

Further, global reactivity descriptors such as global hardness (□ = ΔE/2), global softness (σ = 1/□), electronegativity (χ = - (E_HOMO + E_LUMO)/2), chemical potential (μ = -χ), and electrophilicity index (ω = χ²/2□) were computed. These descriptors offer insights into molecular stability, chemical reactivity, and the potential for interaction with biological targets (Rijal *et al*., 2023). Frontier Molecular Orbital (FMO) analysis, particularly focusing on the HOMO-LUMO gap, was used to assess molecular reactivity and stability. A lower ΔE value indicates higher chemical reactivity, while a larger ΔE gap suggests greater molecular stability. These analyses provide a detailed understanding of the compounds’ potential as drug candidates.

### 2.6. *In silico* ADME/T and toxicity analysis

QikProp Module of Schrödinger, SwissADME (Daina *et al*., 2017), ProTox-3.0 (Banerjee *et al*., 2024), ADMETlab 3.0 (Fu *et al*., 2024), and pkCSM (Pires *et al*., 2015) server were used in the analysis of pharmacokinetic properties to assess the detailed ADMET properties of the two best-ranked compounds based on lowest binding energy from MM-GBSA screening. SwissADME, pkCSM and ProTox-3.0 are free web tools for predicting pharmacokinetics, drug-likeness and medicinal chemistry friendliness of small molecules.

### 2.7. Molecular Dynamics (MD) simulation studies

The MCL1-CNP0161565 and MCL1-CNP0405001 complexes were simulated using GROMACS (2024.2) (Abraham *et al*., 2024) with the CHARMM36 (Brooks *et al*., 1983) force field for parameterization and solvation involving the 3-site TIP3P rigid water model (Mark & Nilsson, 2001). Neutralization was achieved through the addition of Na[and Cl[ions. Further, the complex underwent relaxation through a 20,000-step energy minimization procedure. Subsequently, equilibration was performed, maintaining constant volume (NVT) and pressure (NPT) conditions throughout the MD simulation using the Nose-Hoover thermostat (Evans & Holian, 1985) and the Parrinello–Rahman barostat (Parrinello & Rahman, 1981).

Following this, a production step for the complexes was conducted at a temperature of 310.15 K and a pressure of 1 atm for a duration of 100 ns to assess the system’s stability within an aqueous environment (Elhady *et al*., 2021; Hess *et al*., 2008; Pan *et al*., 2021). Various structural and stability analyses were performed on the MD trajectories. The root mean square deviation (RMSD) was calculated to assess the structural stability of the protein-ligand complex over time using the formula: 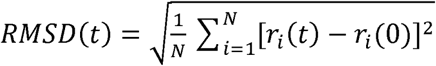 where *r_i_* (*t*) is the position of atom *i* at time *t*, *r_i_*(0) is the initial position, and *N* is the total number of atoms. The root mean square fluctuation (RMSF) was used to evaluate the flexibility of individual residues in the protein during the simulation, calculated as: 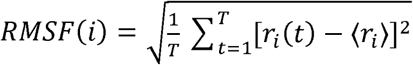, where ⟨*r_i_*⟩ is the position of atom *i* over time. The number of hydrogen bonds (H-bonds) formed between the protein and ligand was monitored throughout the simulation to assess binding stability, as H-bonds are critical for maintaining protein-ligand interactions. The radius of gyration (Rg) was calculated to examine the compactness the protein structure over time, defined as: 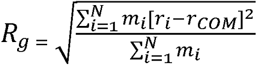 where *m_1_* is the mass of atom *i*. *r_i_* is the position of atom *i* and *r*_COM_ is the centre of mass of the protein. A stable Rg value indicates a well-folded structure. The solvent-accessible surface area (SASA) was computed to analyze the exposure of the protein to the solvent during the simulation. SASA is a measure of the surface area of a molecule that is accessible to a solvent and is critical in understanding protein stability and folding. All the graphs were created in xmgrace. Principal component analysis (PCA) and the free energy landscape (FEL) assessment were conducted by computing and diagonalizing the covariance matrix.

The FEL was formulated as: *F*(*X*) = -*k_B_T* *ln P*(*X*), where *F(X)* represents the free energy, *k_B_* is the Boltzmann constant, *T* is the absolute temperature, and *P(X)* represents the probability distribution of the molecular system along the principal components (David & Jacobs, 2014; Fatima *et al*., 2020). The VMD analysis scripts were used to visualize trajectory files of the MD simulation. Radius of Gyration and RMSD was used to generate the 2D and 3D FEL plots. Python scripts were used for this purpose with Matplotlib package.

## 3. RESULTS

### 3.1. The MCL1 Protein

The X-ray crystallographic structure of the MCL1 target protein [PDB ID: 6O6F], found at https://www.rcsb.org, in complex with inhibitor AMG-176 (*Tapotoclax*) and determined to a resolution of 1.60Å was used in the study (Caenepeel *et al*., 2018).

As illustrated in Figure 1, the protein structure consists of the BH1 domain, highlighted in green, is known to play a crucial role in protein-protein interactions, the BH2 domain, shown in blue, is involved in dimerization, and the BH3 domain, coloured red, is the active site and is responsible for binding to anti-apoptotic proteins.

**Figure 1.**
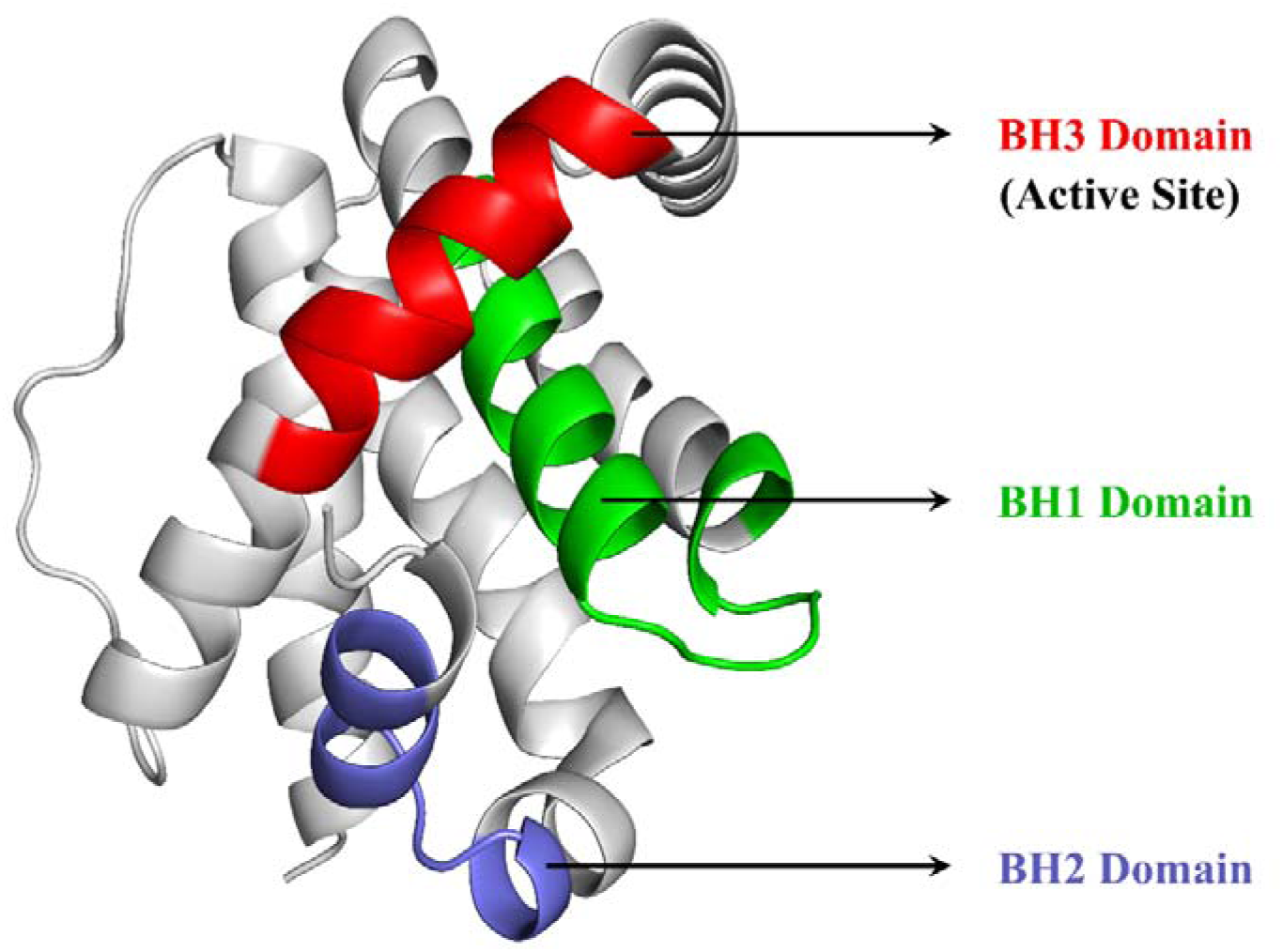
Crystallographic structure of the MCL1 protein with the BH1, BH2, and BH3 domains highlighted in green, blue, and red, respectively.

Analysis through PDBsum web server identified 9 α-helices, 2 γ-turns, 1 β-turns as well as 24 helix-helix interactions. Furthermore, the Ramachandran plot was also used to validate BCL2, and it indicated that 92.6% of the residues were in preferred regions, 7% were in additional residue regions, 0.4% in generous regions, and 0.0% in disallowed regions with a total G- Factor of 0.49. (Ramachandran *et. al.,* 1963) (supplementary figure 1)

### 3.2. Pharmacophore modelling/screening

#### 3.2.1. Diverse ligand based pharmacophore

The generated pharmacophore models were ranked automatically based on the PhaseHypo Score. The 40 generated pharmacophore models with their scoring metrices are listed in supplementary table 1. The hypothesis AAAHHRR_1 revealed the highest PhaseHypo Score of 1.495728 comprising three hydrogen bond acceptor (A), two aromatic rings (R), and two Hydrophobic (H) features as shown in figure 2A & B. The survival score of the hypothesis is 8.263797, site score is 0.719794, vector score is 0.928613, volume score is 0.638179, selectivity score is 3.431904, and BEDROC score is 0.999919.

**Figure 2.**
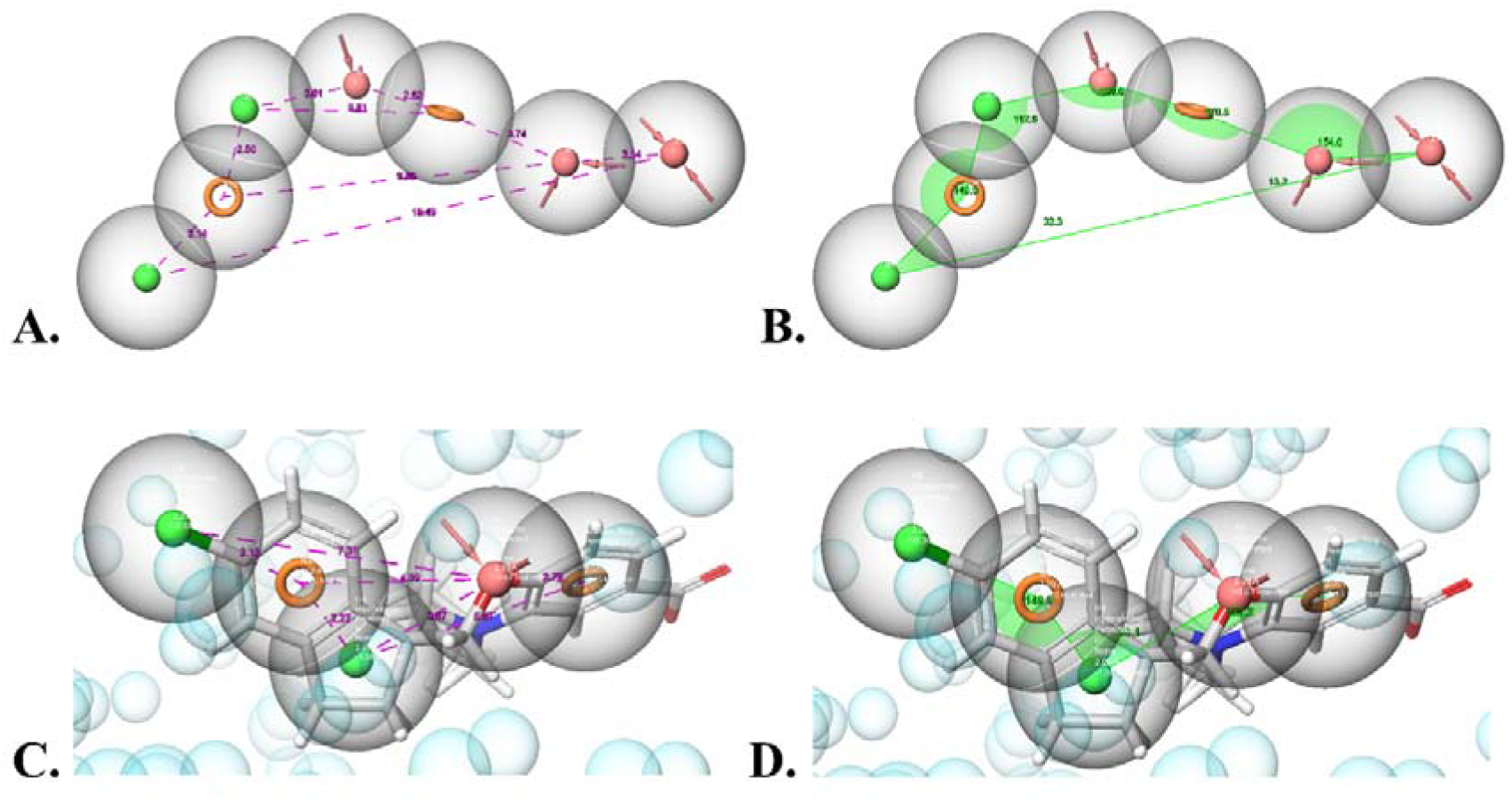
The distance (A) and angles (B) between pharmacophore features of the best ligand-based pharmacophore model, AAAHHRR_1, and the e-pharmacophore model in C and D respectively.

The ∼3.5 million prepared ligand structures were screened using this AAAHHRR_1 pharmacophore model and screened to 4,325 structures.

#### 3.2.2. E-Pharmacophore

The generated e-pharmacophore model consists of four features: one hydrogen bond acceptor (A), two hydrophobic group (H), and two aromatic rings (R), designated as AHHRR. The planar representation of the AHHRR pharmacophore is shown in Figure 2C & D.

#### 3.2.3. Pharmacophore models validation

The pharmacophore model developed was rigorously validated to assess its ability to accurately predict the activity of novel compounds identified through database screening or synthesized de novo. Validation is an essential step before utilizing a pharmacophore model for virtual screening (Kaserer et al., 2015).

The diver ligand based pharmacophore model AAAHHRR_1 model was applied to a test set database comprising 100 inactive molecule generated via the DUDE Decoys tool. It screened the decoy set to only 0 structures indicating an efficiency of ∼100%. Further, validation of the AAAHHRR_1 hypothesis revealed that EF in the top 1% of the decoy dataset is 11.42%, demonstrating that pharmacophore model is 11.42-fold efficient in detecting true positives/actives from the entire dataset. ROC score, RIE, and AUAC values were calculated as 0.99, 9.40, and 0.95, respectively, and shown in figure 3A. Thus, AAAHHRR_1 was statistically significant in picking the actives from the decoy dataset. Statistical significance of model was also validated by calculating BEDROC. Contrary to EF, BEDROC seeks to measure the early enrichment of the actives. BEDROC values were calculated at different tuning parameter values (α[=[8.0, α[=[20.0, and α[=[160.9) and found to be 0.993, 0.996, and 1.000, respectively.

**Figure 3.**
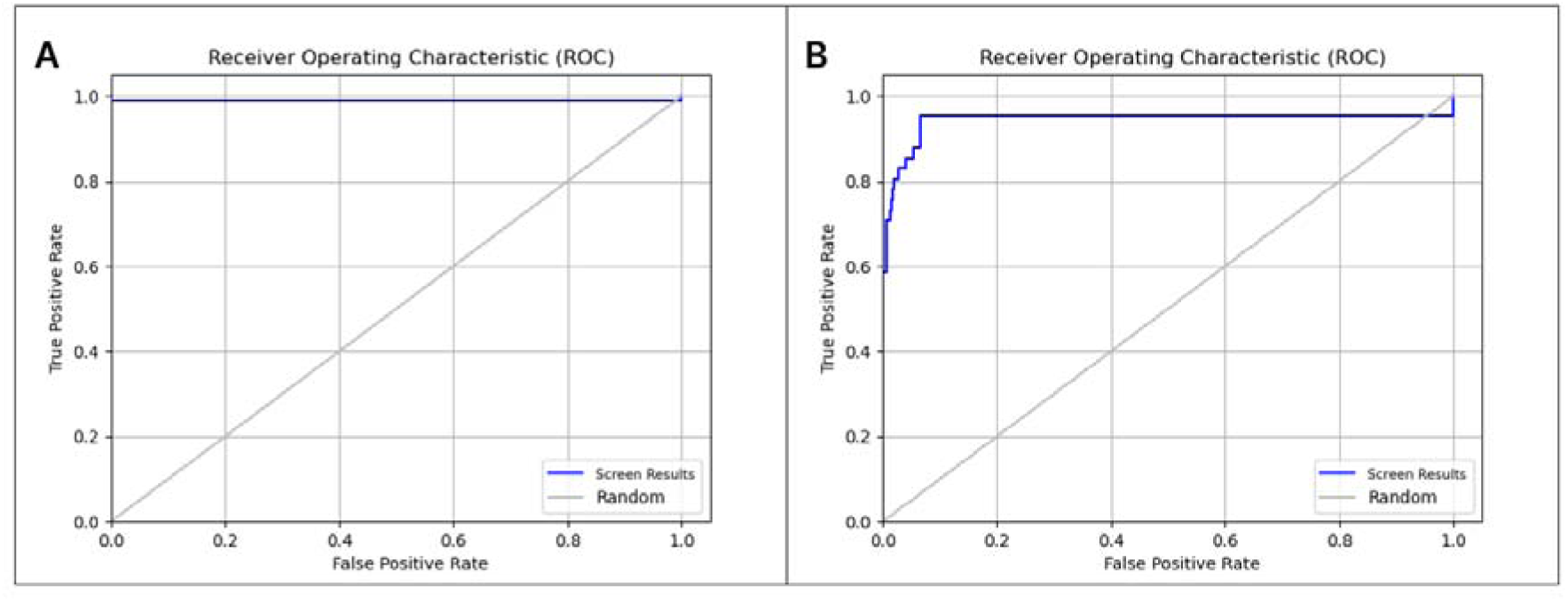
ROC curve represents the pharmacophore validation of the model A: AAAHHRR and B: AHHRR. The solid line represents the performance of the model, while the dashed line represents random guessing. The area under the curve (AUC) provides a measure of overall model performance.

Similarly, the same decoy set was screened using the e-pharmacophore model AHHRR. The e-pharmacophore model was also efficient, screening the decoy set down to 8 structures, achieving an efficiency of ∼92%. The model demonstrated an enrichment factor (EF) of 15.93% in the top 1% of the decoy dataset, indicating it was 15.93 times more effective in identifying true positives. The ROC score, RIE, and AUAC values were 0.79, 8.60, and 0.89, respectively, confirming the model’s statistical significance, depicted in figure 3B. BEDROC values, which assess early enrichment, were 0.889, 0.902, and 0.998 at different tuning parameters (α = 8.0, 20.0, 160.9), further validating the model’s effectiveness.

### 3.3. QSAR Analysis and screening

The top 10 QSAR models were retained and listed in supplementary table 2. The best-fitted model, Kernel Partial Least Squares (KPLS)_Dendritic, exhibiting well-acceptable R^2^ of 0.8997, with S.D. of 0.0022, RMSE of 0.0021 and Q^2^ of 0.8977 was used to predict the IC_50_ of the screened compounds and screen them further (Figure 4). The IC_50_ values of the 2024 screened compounds, observed in the study, ranged from 0.0283 nM to 0.0495 nM, with mean and median values of 0.0302 nM and 0.0372 nM respectively.

**Figure 4.**
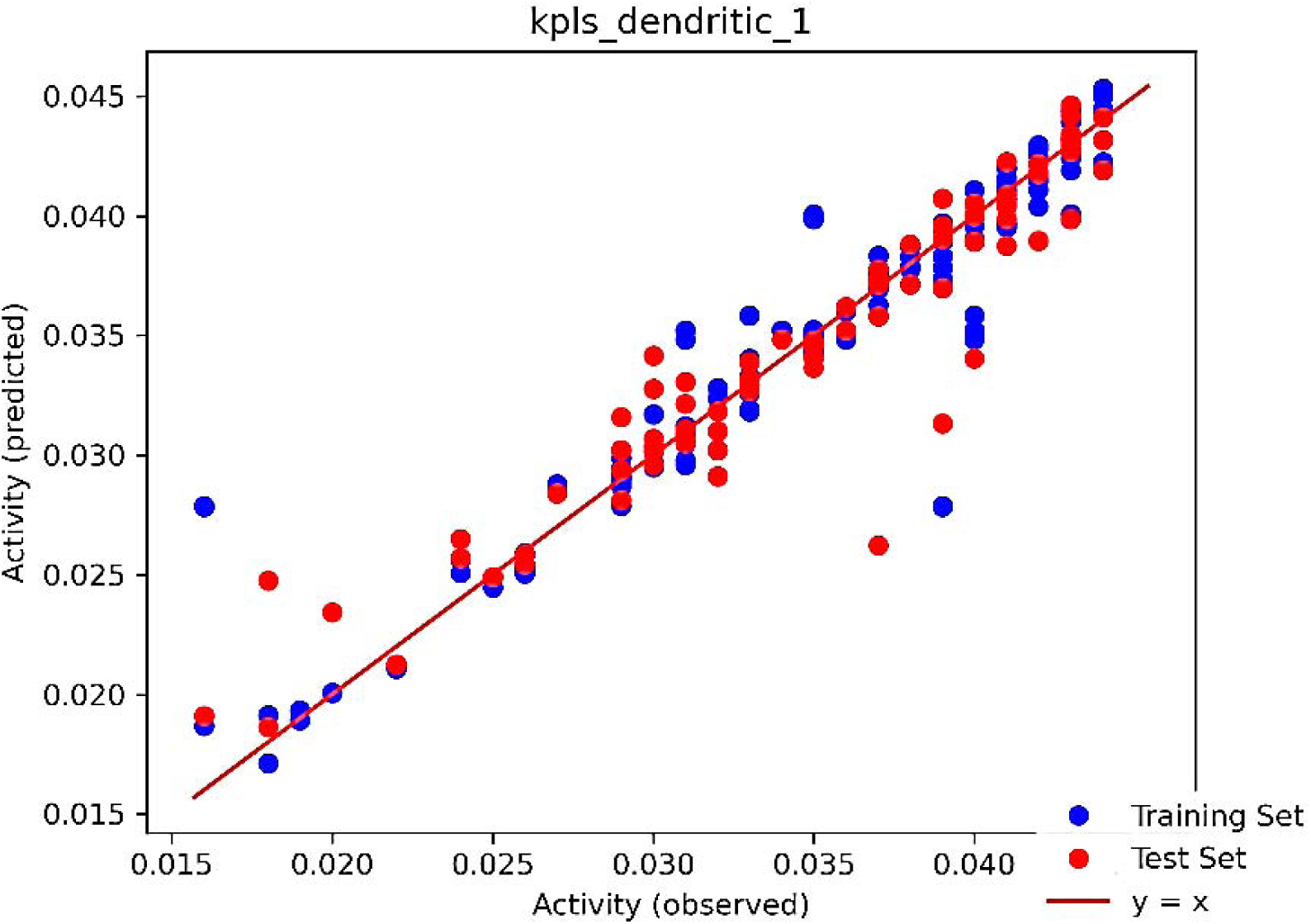
The QSAR model scatter plot of the best model kpls_dendritic_1.

### 3.4. Binding Affinity Prediction Analysis

The docking score range for the top 100 hits compounds was found between -11.793 kcal/mol to -8.232 kcal/mol, after XP docking.

The molecular interactions between the MCL1 protein and the ligands having the best binding affinities - CNP0161565 and CNP0405001 were explored (Figure 5). The results revealed a favourable binding pose characterized by a variety of interactions as listed in the table 1 with the major interacting residues and the interaction distances.

**Figure 5.**
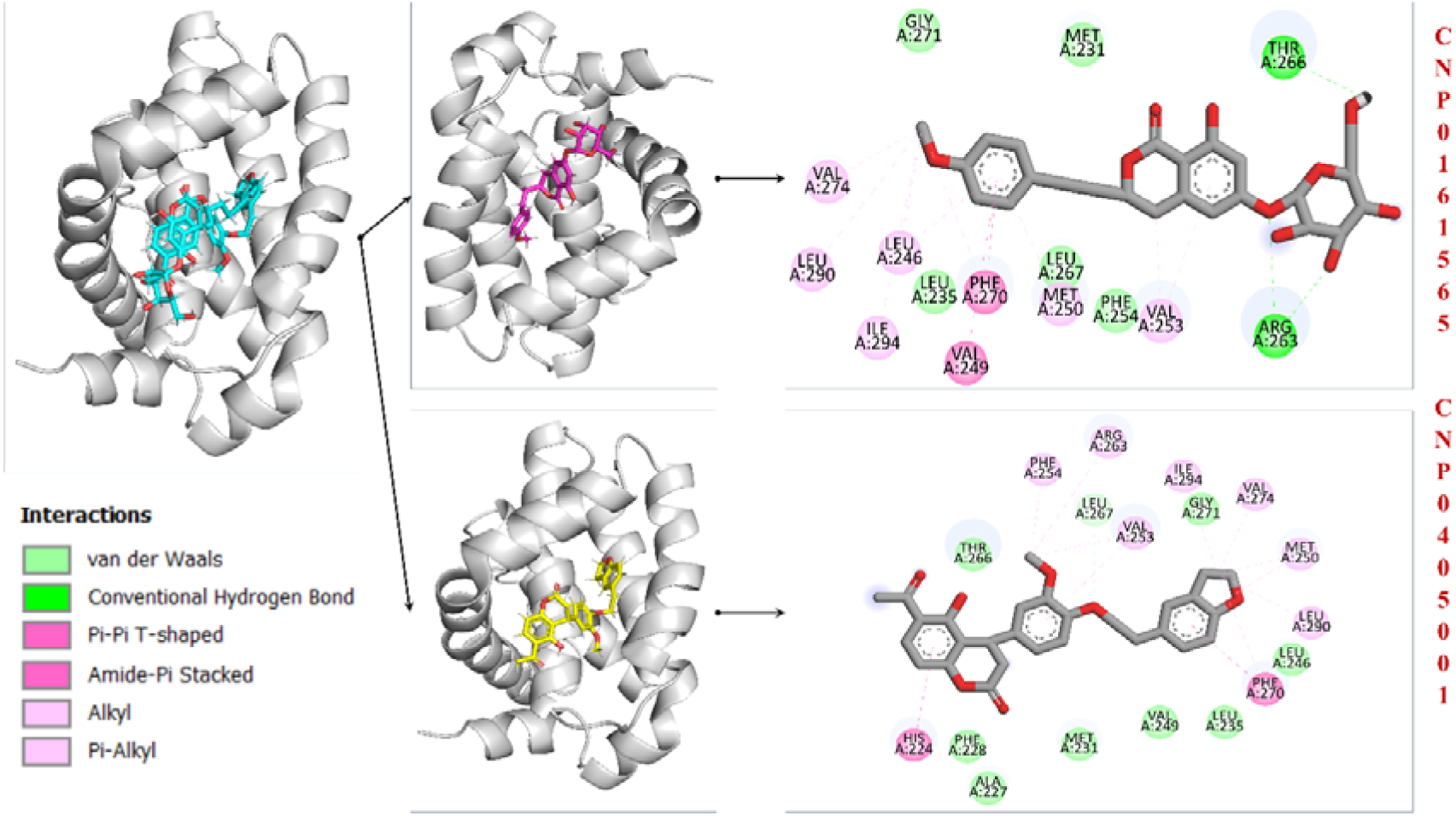
Molecular interactions between the ligands CNP0161565 and CNP0405001 and the MCL1 protein. The figure shows a 3D representation of the protein-ligand complex, and the corresponding 2D interaction highlighting various types of interactions. Key residues involved in these interactions are labelled.

**Table 1.** Molecular interactions and binding affinity of the top 5 ligands based on MM-GBSA score, their COCONUT ID, chemical formula, binding affinities (docking and MM-GBSA), interacting residues and distances.

| COCONUT ID | Formula | Binding Affinity |  | H-Bond Interactions |  |  | Hydrophobic Interactions |
| --- | --- | --- | --- | --- | --- | --- | --- |
|  |  | Docking | MM-GBSA | Donor | Acceptor | Distance |  |
| CNP0161565 | C<sub>24</sub>H<sub>28</sub>O<sub>10</sub> | -10.267 kcal/mol | -79.52 kcal/mol | Arg263:HH12 | L:O3 | 3.075 Å | Phe270 (4.725 Å), Val249 – Met250 (5.248 Å), Val253 (4.813 Å), Leu246 (4.654 Å), Val274 (3.711 Å), Leu290 (4.247 Å), Ile294 (5.307 Å), Phe270 (5.452 Å), Val253 (5.122 Å), Leu246 (5.351 Å), Met250 (4.203 Å) |
|  |  |  |  | Arg263:HH11 | L:07 | 1.853 Å |  |
|  |  |  |  | L:H9 | Thr266:OG1 | 1.978 Å |  |
| CNP0405001 | C<sub>28</sub>H<sub>26</sub>O<sub>7</sub> | -9.962 kcal/mol | -74.68 kcal/mol | Leu267:HA | L:05 | 2.711 Å | His224 (4.812 Å), Phe270 (4.846 Å), Met250 (4.486 Å), Val274 (4.871 Å), Leu290 (5.419 Å), Ile294 (4.883 Å), Val253 (5.427 Å), Arg263 (4.089 Å), Phe254 (5.302 Å), Phe270 (5.085 Å), Val253 (4.870 Å), Met250 (4.119 Å) |
| CNP0353199 | C<sub>27</sub>H<sub>26</sub>O<sub>7</sub> | -9.071 kcal/mol | -72.87 kcal/mol | Leu267:HA | L:05 | 2.678 Å | His224 (4.947 Å), Phe270 (4.839 Å), Met250 (4.449 Å), Val274 (3.310 Å), Leu290 (4.344 Å), Ile294 (5.468 Å), Val253 (5.437 Å), |
|  |  |  |  |  |  |  | Arg263 (4.156 Å), Phe254 (5.449 Å), Phe270 (5.402 Å), Val253 (4.939 Å), Met250 (4.093 Å) |
| CNP0420896 | C<sub>26</sub>H<sub>26</sub>O<sub>8</sub> | -9.879 kcal/mol | -72.68 kcal/mol | L:H3 | His224:ND1 | 1.885 Å | Ala227 (3.447 Å), Phe270 (3.786 Å), His224 (3.817 Å), Met250 (4.138 Å), Ile294 (4.158 Å), Val249 (4.209 Å), Met231 (4.237 Å), Met250 (4.304 Å), Ala227 (4.363 Å), Met231 (4.434 Å), Leu235 (4.465 Å), Phe270 (4.543 Å), Phe228 (4.817 Å), Phe270 (4.821 Å), Val253 (4.838 Å), Leu267 (4.920 Å), Val253 (5.244 Å), Leu267 (5.270 Å) |
|  |  |  |  | L:H12 | Leu267:O | 2.089 Å |  |
| CNP0369034 | C<sub>24</sub>H<sub>24</sub>O<sub>5</sub> | -9.122 kcal/mol | -72.57 kcal/mol | - |  |  | Met250 (3.984 Å), His224 (4.043 Å), Met250 (4.362 Å), Phe270 (4.798 Å), Phe270 (4.928 Å), Val249 – Met250 (4.930 Å), Phe270 (5.081 Å), Leu246 (5.138 Å), Val253 (5.202 Å), Val274 (5.337 Å), Ala227 (5.341 Å), Ile294 (5.377 Å), His224 (5.730 Å) |

### 3.5. Density Functional Theory (DFT) Studies

#### 3.5.1. Geometry optimization and FMO (HOMO/LUMO) analysis

The highest occupied molecular orbital (HOMO) defines the ability of a molecule to donate electrons. The lowest unoccupied molecular orbital (LUMO) tells about its capacity to accept electrons. These orbitals are important in describing the interaction of a molecule with another species, the molecular reactivity, electronic structure, kinetic stability, and hardness or softness. Besides, the energy difference between HOMO and LUMO is very important for the estimation of chemical reactivity. The larger the energy gap, the higher the stability and lower the reactivity. The smaller it is, the poorer the stability and the greater the reactivity. Depending on the basis set applied, our research demonstrated compounds CNP0405001 and CNP0161565 as having the smallest value for the energy gap having ΔE as 0.11351 kcal/mol and 0.12036 kcal/mol respectively – they are the least stable and most reactive, according to Figure 6. The reactivity of the compounds increases from the lowest to the highest order: CNP0405001 < CNP0161565 < CNP0369034 < CNP0353199 < CNP0420896. Geometry optimization was performed by using FMO-based DFT techniques, and the calculation of the HOMO-LUMO energy band was done by using the Gaussian software, which was aimed to predict the reactivity of the above-mentioned compounds.

**Figure 6.**
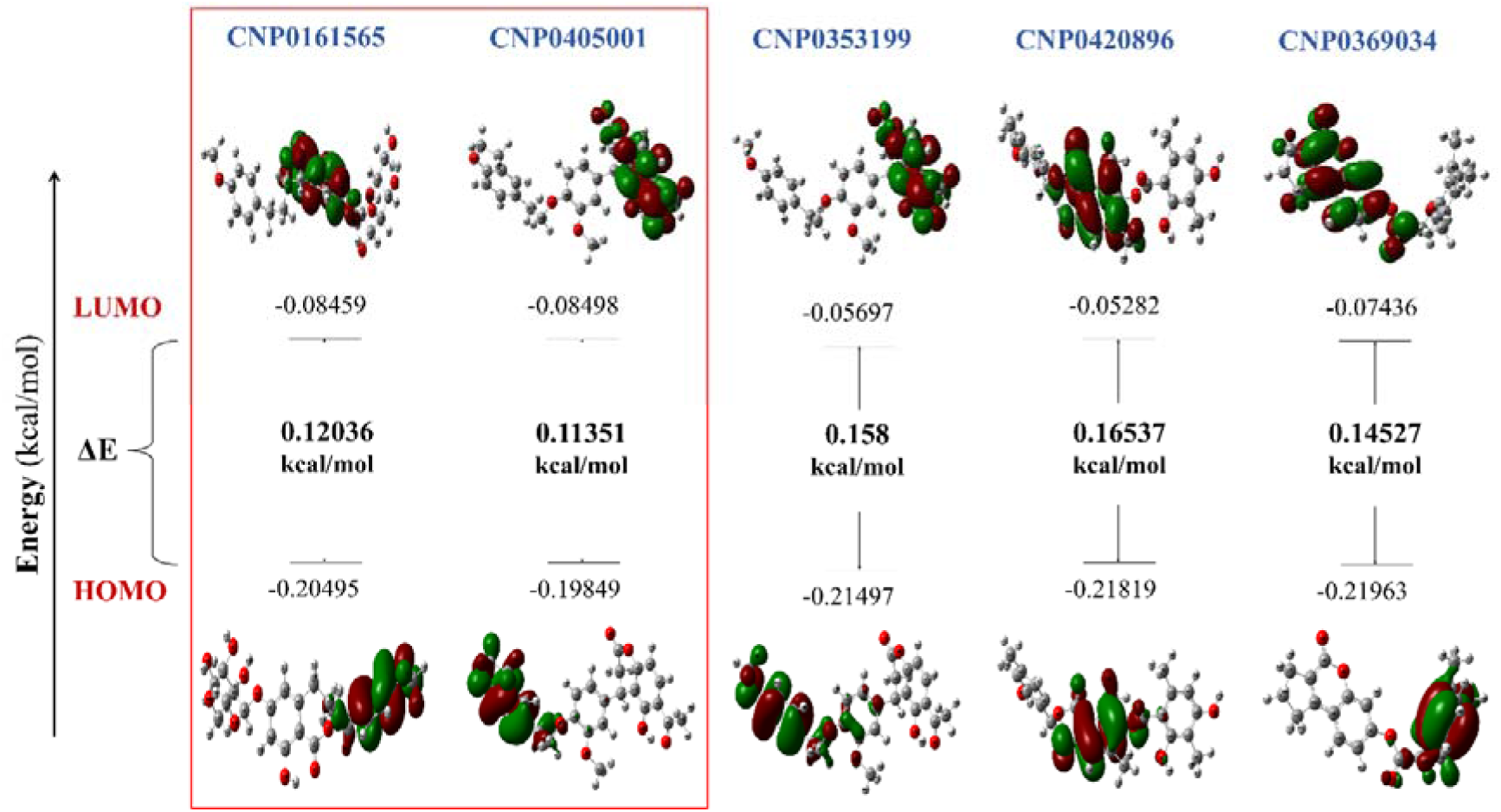
Frontier molecular orbitals (FMOs) illustrating the HOMO and LUMO electron density distributions of five ligands, along with their corresponding energy values of the top 5 ligands.

#### 3.5.2. Quantum chemical calculations

The reactivity and stability of the compounds can be analyzed using a range of quantum chemical parameters, including ionization potential (IP), electron affinity (EA), hardness (η), softness (S), chemical potential (μ), electrophilicity index (ω), electronegativity (χ), total energy, and dipole moment. The ionization potential (IP), which reflects the energy required to remove an electron, is derived from the negative of the HOMO energy, while electron affinity (EA), indicating the energy gained when an electron is added, is calculated from the negative of the LUMO energy. For CNP0161565, the IP is 0.04726 kcal/mol and the EA is 0.02013 kcal/mol. CNP0405001 has an IP of 0.04519 kcal/mol and an EA of 0.02028 kcal/mol, while for CNP0353199, the IP and EA are 0.04941 kcal/mol and 0.01321 kcal/mol, respectively. Similarly, CNP0420896 exhibits an IP of 0.05021 kcal/mol and an EA of 0.01106 kcal/mol, and CNP0369034 has values of 0.05052 kcal/mol for IP and 0.01806 kcal/mol for EA. The quantum chemical parameters for the compounds are summarized in Table 2.

**Table 2.** Quantum molecular descriptors of the top 5 ligands.

| **Compound** | **IP** | **EA** | **$\eta$** | **S** | **$\mu$** | **$\omega$** | **$\chi$** | **Energy** | **Dipole Moment** |
| --- | --- | --- | --- | --- | --- | --- | --- | --- | --- |
| CNP0161565 | -0.20495 | -0.08459 | 0.06018 | 16.61957 | -0.14477 | 6.35527 | 0.14477 | -0.00001254 | 10.809 |
| CNP0405001 | -0.19849 | -0.08498 | 0.05675 | 17.60136 | -0.14174 | 6.30445 | 0.14174 | -0.00001228 | 6.680 |
| CNP0353199 | -0.21497 | -0.05697 | 0.07900 | 12.65822 | -0.13597 | 6.17425 | 0.13597 | -0.00001179 | 7.453 |
| CNP0420896 | -0.21819 | -0.05282 | 0.08269 | 12.07330 | -0.13551 | 6.08239 | 0.13551 | -0.00001172 | 9.153 |
| CNP0369034 | -0.21963 | -0.07436 | 0.07264 | 13.77006 | -0.14700 | 6.45358 | 0.14700 | -0.00001274 | 8.009 |

The hardness (η), which indicates the resistance to electron cloud deformation, is calculated as half the difference between the IP and EA. The hardness values are as follows: CNP0161565 (0.03032 kcal/mol), CNP0405001 (0.02893 kcal/mol), CNP0353199 (0.03610 kcal/mol), CNP0420896 (0.03908 kcal/mol), and CNP0369034 (0.03673 kcal/mol). Softness (S), which is the inverse of hardness, reveals that CNP0405001 is the softest, indicating higher reactivity, while CNP0420896 is the hardest, implying greater stability.

Chemical potential (μ), the negative average of the HOMO and LUMO energies, provides insight into the likelihood of a compound gaining or losing electrons. CNP0161565 and CNP0405001 have chemical potentials of -0.07212 kcal/mol and -0.07089 kcal/mol, respectively, both indicating good stability and a strong ability to gain electrons. Other compounds, such as CNP0353199 (-0.06720 kcal/mol), CNP0420896 (-0.06703 kcal/mol), and CNP0369034 (-0.07373 kcal/mol), also exhibit reasonable chemical potential values, but CNP0161565 and CNP0405001 stand out as more stable.

The electrophilicity index (ω), calculated as μ²/2η, reflects a molecule’s ability to accept electrons. Higher values suggest greater electron acceptance capacity. In this case, CNP0161565 and CNP0405001 have moderate electrophilicity values, reinforcing their balance between reactivity and stability. Electronegativity (χ), the negative of chemical potential, shows that CNP0161565 and CNP0405001 have values of 0.07212 kcal/mol and 0.07089 kcal/mol, respectively, indicating a favorable tendency to attract electrons.

The total energy, measured in atomic units (a.u.), further supports the stability of these compounds, while dipole moment values, which reflect molecular polarity and binding affinity, confirm their potential as drug candidates. CNP0161565 exhibits a dipole moment of 10.809 Debye, while CNP0405001 shows a dipole moment of 6.680 Debye, indicating excellent binding potential.

Based on the quantum chemical parameters calculated, CNP0161565 and CNP0405001 are the most promising candidates for drug development. Their combination of favorable chemical potential, optimal hardness, balanced electrophilicity, and strong dipole moments make them highly suitable for further investigation as potential therapeutic agents.

### 3.6. ADME/T Analysis

Both CNP0161565 and CNP0405001 exhibit favourable drug-like properties, adhering to Lipinski’s Rule of Five and possessing balanced physicochemical characteristics as depicted in figure 7A & B. The physiochemical, medicinal chemistry and ADMET properties of CNP0161565 and CNP0405001 are summarized in details in the supplementary figures 3, 4 & 5. While CNP0161565 demonstrates a more complex structural architecture with multiple rings and heteroatoms, CNP0405001 presents a simpler structure. Both compounds exhibit moderate polarity and hydrophobicity, which can influence their solubility and membrane permeability. CNP0161565 displays a better balance of hydrogen bond donors and acceptors, potentially enhancing its interaction with biological targets.

**Figure 7.**
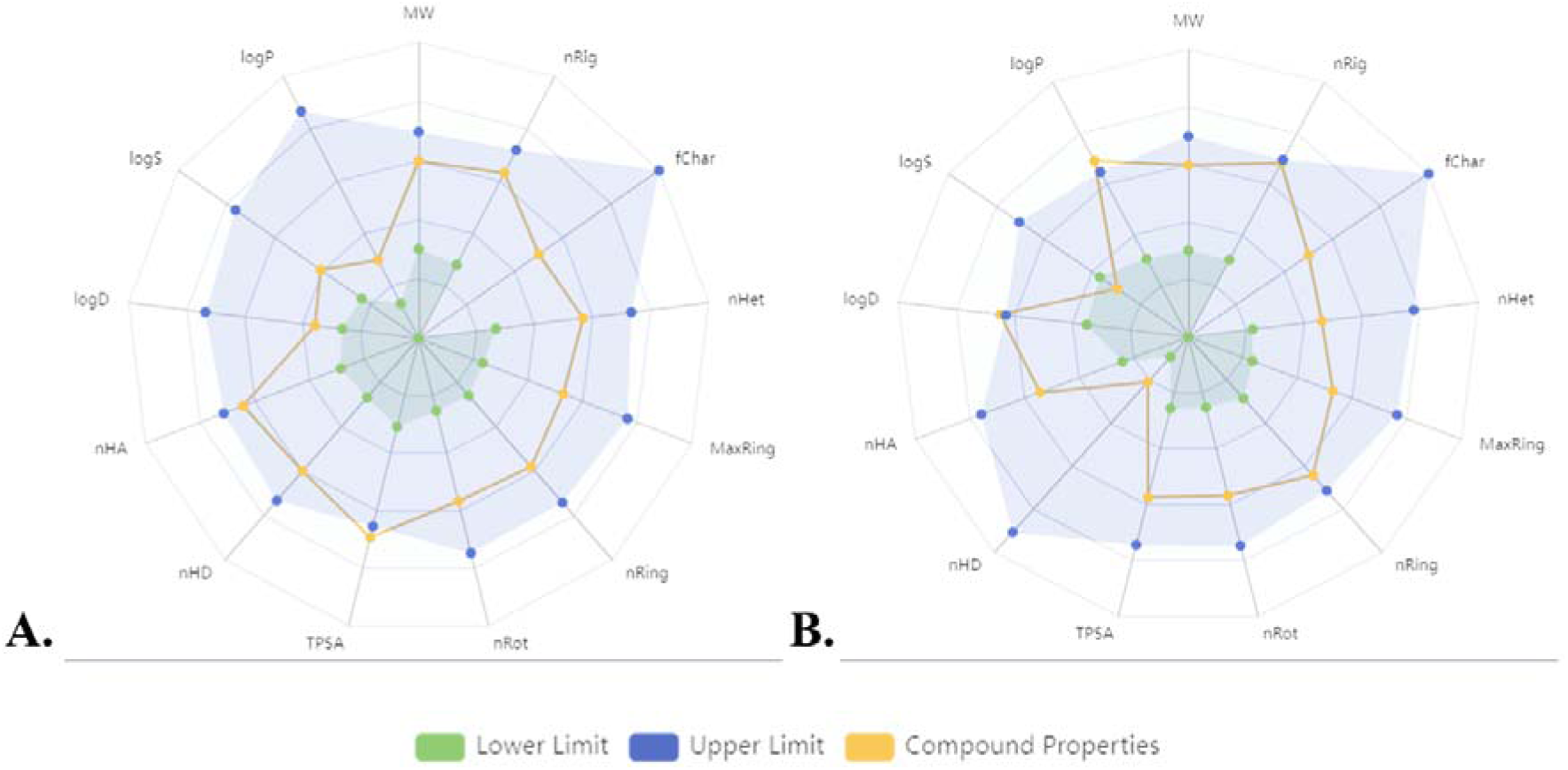
The bioavailability radar plot (obtained from ADMETlab 3.0) depicting the excellent drug-likeliness of the two compounds – A: CNP0161565; B: CNP0405001.The blue area represents the optimal range for each property, and the yellow dots represents the properties of the compounds.

Regarding solubility, CNP0161565 and CNP0405001 both suffer from poor water solubility, which might necessitate formulation strategies to enhance oral bioavailability (supplementary figure 2). However, CNP0405001 demonstrates superior human intestinal and oral absorption compared to CNP0161565. Both compounds face challenges in terms of membrane permeability, as evidenced by low Caco-2 and MDCK permeability values. While CNP0161565 is a substrate for P-glycoprotein, CNP0405001 inhibits this efflux transporter, which could potentially impact the bioavailability of co-administered drugs.

In terms of distribution, CNP0161565 exhibits moderate distribution with significant plasma protein binding, whereas CNP0405001 displays limited tissue distribution due to high plasma protein binding. Both compounds demonstrate poor CNS penetration, reducing the risk of central nervous system-related side effects. The boiled egg plot depicting the GI absorption and BBB penetration of two compounds is shown in supplementary figure 3. Metabolically, CNP0161565 demonstrates a cleaner profile with no major CYP enzyme interactions, while CNP0405001 is a substrate and inhibitor of multiple CYP enzymes, increasing the potential for drug-drug interactions. Excretion appears to be primarily non-renal for both compounds. Toxicologically, CNP0161565 and CNP0405001 exhibit acceptable safety profiles with low acute toxicity and no significant genotoxicity or carcinogenicity concerns. However, CNP0405001 raises a potential cardiotoxicity flag due to hERG inhibition, warranting further investigation. Both compounds show promise as drug candidates but require optimization in areas such as solubility, permeability, and metabolic liabilities. Additionally, formulation strategies to address bioavailability and distribution challenges are crucial.

Overall, both compounds present interesting pharmacological profiles with potential for further development. A careful balance of optimizing their physicochemical properties, addressing ADME limitations, and mitigating potential toxicities will be essential in advancing these compounds towards clinical candidates.

### 3.7. Molecular Dynamics Simulation

#### 3.7.1. RMSD analysis

##### 3.7.1.1. Protein RMSD

The RMSD (Root Mean Square Deviation) analysis for the MCL1 protein bound to inhibitors CNP0161565 (black curve) and CNP0405001 (red curve) reveals insights into structural stability during simulations, shown in figure 8A. Initially, both complexes show RMSD values around 1 Å, indicating good equilibration and stable structural integrity, suggesting minimal conformational changes upon binding.

**Figure 8.**
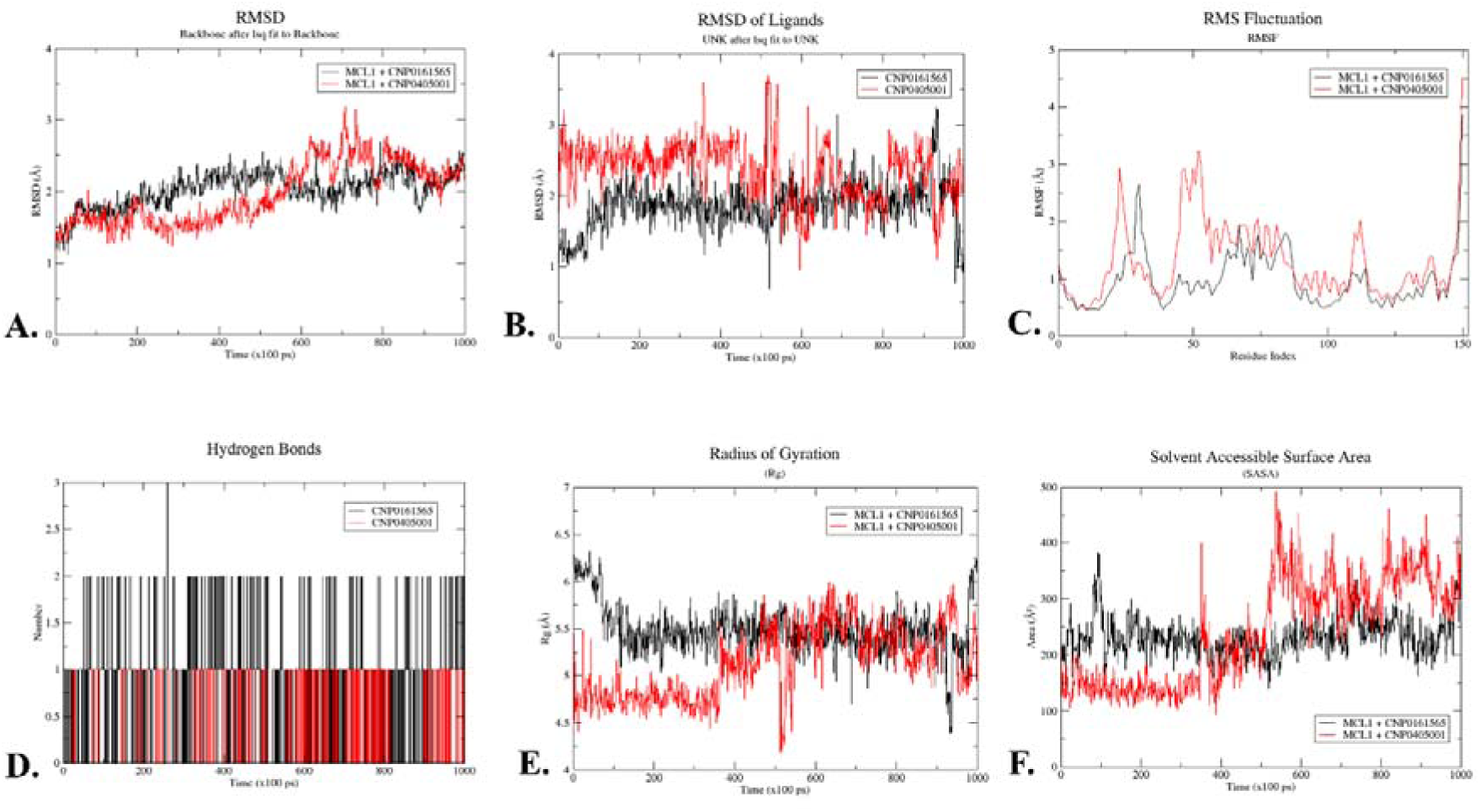
Time-dependent changes in RMSD, RMSF, H-Bond formation, RoG, and SASA for two ligand-protein complexes – MCL1- CNP0161565 (in black) and MCL1- CNP0405001 (in red)

As the simulation progresses, the MCL1-CNP0161565 complex’s RMSD gradually increases, stabilizing around 2 Å, reflecting minor conformational adjustments and a stable binding mode that supports the native protein conformation—essential for inhibitor efficacy.

Conversely, the MCL1-CNP0405001 complex exhibits a more dynamic RMSD profile, initially rising to around 2 Å before peaking at approximately 3.2 Å at the 600 ns mark, indicating significant conformational changes or destabilization. After this peak, the RMSD oscillates around 2.5 Å, suggesting that CNP0405001 induces greater structural flexibility in MCL1, potentially leading to multiple binding modes or transient unfolding.

Comparatively, CNP0161565 provides a stable interaction with MCL1, maintaining a rigid conformation beneficial for therapeutic contexts requiring structural integrity. In contrast, the higher RMSD and flexibility of MCL1-CNP0405001 imply a different mode of action, allowing for larger conformational changes that may be advantageous for dynamic interactions but could result in less stable binding and variable efficacy.

This analysis highlights the distinct interactions of the two inhibitors: CNP0161565 stabilizes MCL1, while CNP0405001 promotes greater flexibility. Understanding these dynamics is essential for optimizing inhibitor efficacy and tailoring therapeutic approaches to specific biological contexts.

##### 3.7.1.2. Ligand RMSD

The RMSD analysis of ligands CNP0161565 (black curve) and CNP0405001 (red curve) within the MCL1 protein reveals their stability and conformational dynamics during the simulation (figure 8B). Both ligands start with RMSD values around 1 Å, indicating well-equilibrated binding within MCL1, suggesting stable binding with minimal initial deviations.

As the simulation progresses, distinct behaviors emerge: CNP0161565 exhibits a stable RMSD trajectory, fluctuating between 1.5 Å and 2.5 Å, indicating strong interactions and consistent binding conformation. In contrast, CNP0405001 displays a more dynamic profile, with RMSD values ranging from 1.5 Å to 3 Å and occasional spikes (up to 3.2 Å) at 300 ns and 600 ns, suggesting transient destabilizations and potential partial dissociations.

Comparatively, CNP0161565 maintains a strong, stable interaction with MCL1, which may enhance its therapeutic efficacy. Conversely, CNP0405001’s fluctuating RMSD suggests greater conformational flexibility, which could imply weaker binding stability. This flexibility may allow CNP0405001 to interact with multiple protein conformations, but it could also indicate less specificity.

The RMSD analysis highlights that CNP0161565 shows lower and more stable values, denoting strong binding, while CNP0405001’s higher and more variable values indicate increased flexibility and transient destabilizations. Understanding these dynamics is essential for optimizing ligand interactions and improving inhibitor design tailored to specific biological contexts.

#### 3.7.2. RMSF Analysis

The Root Mean Square Fluctuation (RMSF) analysis provides insights into the flexibility and stability of protein residues during molecular dynamics (MD) simulations. RMSF values indicate how much each residue fluctuates around its average position, reflecting the protein-ligand complex’s dynamic behavior. Lower RMSF values indicate more stable protein regions, while higher values suggest flexibility. This analysis focuses on MCL1 protein in the presence of two ligands—CNP0161565 (black curve) and CNP0405001 (red curve)—to understand their impact on protein behavior, as shown in figure 8C.

For MCL1 + CNP0161565, the black curve shows stable RMSF values across most residues, indicating effective stabilization by the ligand and strong protein interaction. In contrast, the red curve for MCL1 + CNP0405001 demonstrates slightly higher RMSF values but maintains an overall stable profile, suggesting effective interaction and adaptive flexibility that may be advantageous therapeutically.

In the initial residue region (0-30), MCL1 + CNP0161565 shows minimal fluctuation, indicative of a stable binding environment crucial for functional conformation. MCL1 + CNP0405001 shows slightly higher RMSF values, suggesting moderate flexibility that allows necessary conformational changes while retaining stability. In the middle residue region (30- 70), MCL1 + CNP0161565 has moderate RMSF values with peaks around residues 40-50, indicating supported flexibility in loop regions. For MCL1 + CNP0405001, the red curve exhibits higher peaks, suggesting enhanced flexibility that may benefit adaptive interactions in therapeutic contexts. In the core residue region (70-110), MCL1 + CNP0161565 shows lower RMSF values, indicating stability essential for structural integrity. MCL1 + CNP0405001 has slightly higher RMSF values, suggesting retained functionality with dynamic capability for molecular interactions. In the terminal residue region (110-150), MCL1 + CNP0161565 shows rising RMSF values typical of flexible terminal regions, indicating effective anchoring of the protein. MCL1 + CNP0405001 similarly shows increased RMSF, suggesting greater flexibility in terminal regions, potentially beneficial for certain functions.

CNP0161565 excels in stabilizing the MCL1 protein, evidenced by lower RMSF values and essential for proper function, making it a strong candidate as an effective inhibitor. CNP0405001, while slightly more flexible, still maintains crucial stability for therapeutic applications. Its adaptive flexibility may allow broader molecular interactions and necessary conformational changes, suggesting versatility as a drug candidate.

CNP0161565 offers superior stabilization of MCL1, ensuring a consistent conformation, while CNP0405001 presents a balance of stability and flexibility, enhancing adaptability in dynamic environments. Both compounds demonstrate significant potential as drug candidates targeting MCL1, with CNP0161565 being particularly promising for rigid inhibition and CNP0405001 for adaptable therapeutic contexts.

#### 3.7.3. Hydrogen bond analysis

The MCL1 complex with CNP0161565 exhibits fluctuating hydrogen bonds (0-3) throughout the simulation, indicating inconsistent interactions and potential instability of binding.

Notably, there are periods without hydrogen bonds (200 ps - 400 ps and near 800 ps), suggesting the ligand may partially dissociate or adopt unfavorable conformations, thereby compromising its efficacy as an inhibitor. In contrast, the MCL1-CNP0405001 complex shows a more stable hydrogen bond pattern, oscillating mainly between 0 and 2, with fewer instances of complete dissociation. This consistent presence of hydrogen bonds indicates that CNP0405001 maintains closer and more continuous contact with MCL1, enhancing its inhibitory potential through stronger interactions (figure 8D).

The hydrogen bond analysis highlights that stable hydrogen bonds are critical for effective protein-ligand interactions. The sporadic bonding in the MCL1-CNP0161565 complex could lead to weaker binding affinity, while the stable hydrogen bonds in the MCL1-CNP0405001 complex suggest a more favorable interaction profile, allowing for better stabilization of MCL1 and more effective inhibition.

Combining hydrogen bond data with radius of gyration (Rg) analysis reveals that CNP0405001 is the more promising candidate, demonstrating stable hydrogen bonds and a compact protein-ligand complex, whereas CNP0161565’s inconsistent bonding and higher Rg values imply less effective binding and reduced inhibitory potential.

#### 3.7.4. Radius of Gyration (Rg)

The graph in figure 8E shows that the MCL1 complex with CNP0161565 (black curve) has a higher average radius of gyration (Rg) of around 5.5 Å (fluctuating between 5.0 and 6.5 Å), indicating it is less compact and more expanded compared to the MCL1 complex with CNP0405001 (red curve), which has a lower average Rg fluctuating between 4.5 and 5.5 Å. This suggests that the MCL1-CNP0405001 complex is more compact and may indicate a more stable protein-ligand interaction.

Significant initial fluctuations in the black curve, particularly in the first 200 ps (Rg oscillating between 6.0 Å and 6.5 Å), stabilize slightly but remain pronounced compared to the more consistent fluctuations of the red curve (Rg largely within 4.5 Å to 5.5 Å). The fewer fluctuations in the MCL1-CNP0405001 complex imply a more stable binding interaction, which reduces conformational freedom for the protein.

The structural compactness and fluctuation stability directly affect ligand efficacy. The more stable MCL1-CNP0405001 complex suggests that this ligand may induce a favorable conformational state in MCL1, enhancing its inhibitory potential. Conversely, the higher Rg and greater fluctuations in the MCL1-CNP0161565 complex indicate a less stable binding mode, potentially reducing inhibitory effectiveness.

Throughout the simulation, the black curve experiences periods of stability interspersed with sharp Rg changes, indicating transient conformational shifts or unfolding. The red curve, while showing some fluctuations, maintains a lower Rg, suggesting fewer large-scale conformational changes. Notably, around 900 ps, the black curve shows a drop in Rg, indicating late-stage compaction or structural rearrangement, while the red curve remains consistent, supporting that CNP0405001 better stabilizes MCL1.

The comparative analysis of Rg values indicates that CNP0405001 induces a more stable and compact conformational state in MCL1 than CNP0161565, with reduced fluctuations and lower Rg values suggesting stronger binding interactions that enhance its potential as a therapeutic inhibitor. Conversely, the higher Rg and greater fluctuations of CNP0161565 may negatively impact its efficacy as an inhibitor.

#### 3.7.5. Solvent Accessible Surface Area (SASA)

The SASA analysis underscores the importance of stability and flexibility in ligand design, influencing binding efficacy and therapeutic potential. Understanding these dynamics is crucial for optimizing inhibitor interactions with target proteins.

SASA analysis assesses the solvent exposure of protein-ligand complexes, revealing differences in binding stability. The SASA profiles for MCL1 with inhibitors CNP0161565 (black curve) and CNP0405001 (red curve) demonstrate significant variations in solvent exposure and dynamics (Figure 8F).

Initially, both complexes have low SASA values, with CNP0161565 averaging ∼200 Å² and CNP0405001 at ∼150 Å², indicating stable binding conformations with ligands well-buried in the binding pocket. As the simulation progresses, fluctuations in SASA values reflect the dynamic behavior of these complexes. The black curve (CNP0161565) fluctuates between 150 Å² and 300 Å², with peaks up to 350 Å², suggesting moderate solvent exposure due to minor conformational adjustments while largely maintaining stability.

In contrast, the red curve (CNP0405001) exhibits broader fluctuations from 100 Å² to 450 Å², with peaks around 400 ns and 600 ns, indicating significant solvent exposure and greater conformational flexibility. This behavior suggests that CNP0405001 may experience partial unbinding events, reflecting a less stable interaction with MCL1.

Overall, CNP0161565 shows a stable interaction with lower, consistent SASA values, which may enhance drug efficacy through secure binding. Conversely, the higher and variable SASA values for CNP0405001 indicate greater flexibility and less specific interaction with MCL1, potentially impacting its binding affinity and inhibitory efficacy.

#### 3.7.6. Binding Modes Analysis

The dynamic behaviour of the ligands CNP0161565 and CNP0405001 within the MCL1 binding site was further examined by visualizing their conformational changes at five distinct time points during the 100 ns molecular dynamics (MD) simulation, as shown in Figure 9.

**Figure 9.**
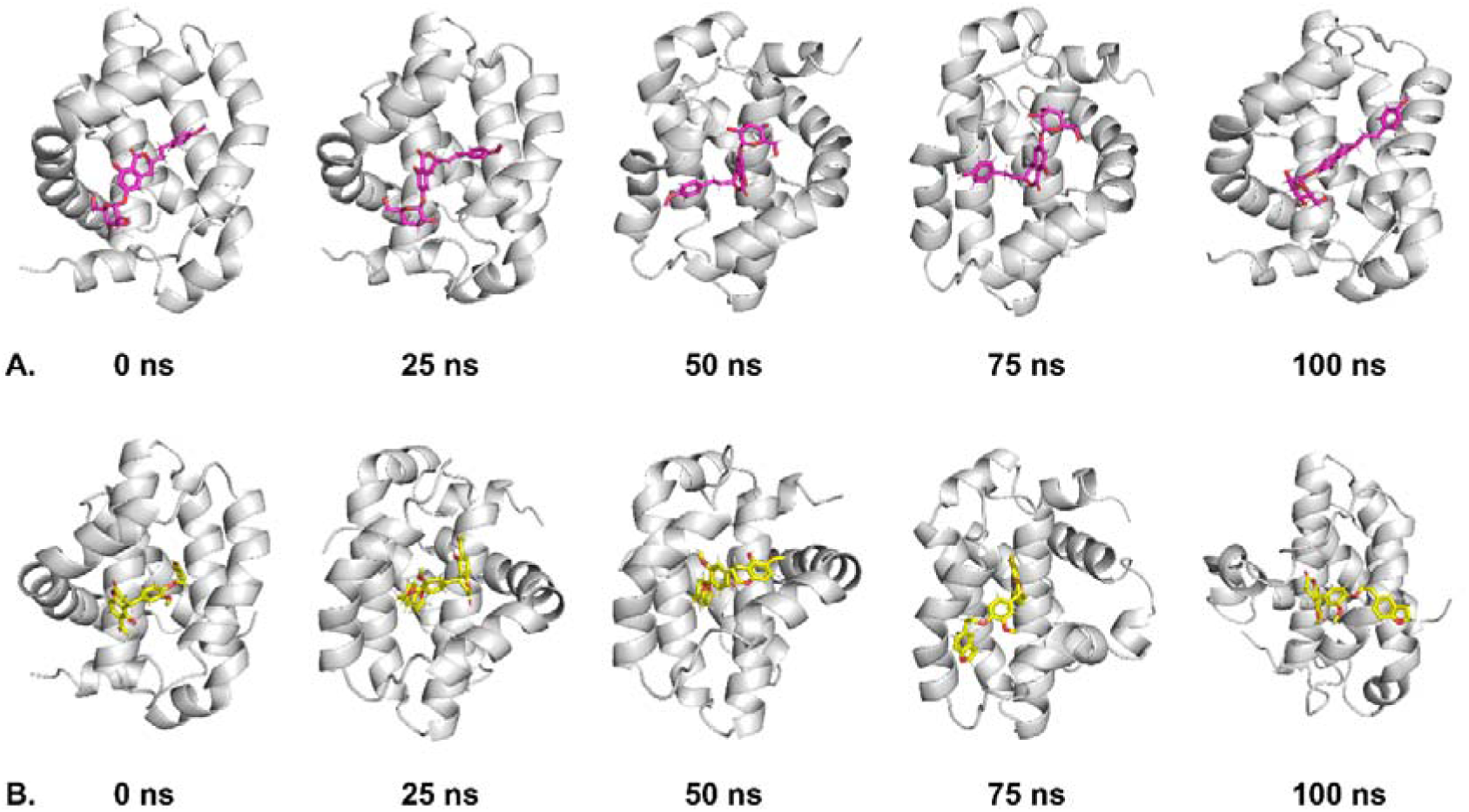
Conformational dynamics of ligands CNP0161565 and CNP0405001 bound to MCL1 over 100 ns of MD simulation. Panel A illustrates the ligand CNP0161565 (purple) and panel B illustrates the ligand CNP0405001 (yellow) bound to the MCL1 protein at five distinct time points: 0 ns, 25 ns, 50 ns, 75 ns, and 100 ns.

This analysis provides insight into the stability and flexibility of each ligand in the binding pocket over time. In panel 9A, the ligand CNP0161565 (represented in purple) shows a relatively stable binding pose throughout the simulation. At all five time points (0 ns, 25 ns, 50 ns, 75 ns, and 100 ns), the ligand maintains its core interactions with the binding pocket of MCL1, with only minor reorientations observed. These minor adjustments suggest that the ligand is optimizing its interactions while maintaining a consistent binding mode, indicating a stable and strong affinity for the target protein. This observation aligns with the results from the RMSD and hydrogen bond analyses, where CNP0161565 demonstrated lower deviations and a more stable hydrogen bonding profile, supporting its high binding stability.

In contrast, panel 9B depicts the behaviour of the ligand CNP0405001 (shown in yellow), which exhibits greater conformational flexibility over the course of the simulation. Notably, as the simulation progresses, especially after 50 ns, the ligand undergoes more significant shifts in its orientation within the binding site. These fluctuations are particularly evident at the 75 ns and 100 ns time points, where the ligand adopts distinct conformations compared to its initial pose. This increased flexibility could indicate weaker interactions with MCL1 or a more dynamic exploration of multiple binding poses within the active site. Such behaviour is reflected in the higher RMSD values and a reduced number of persistent hydrogen bonds observed for CNP0405001. The structural instability and increased ligand mobility suggest that CNP0405001 has a less favourable binding profile compared to CNP0161565.

#### 3.7.7. Principal Component Analysis (PCA)

The PCA projections depict the conformational sampling of the MCL1 protein with two ligands, A (CNP0161565) and D (CNP0405001) in figure 10. Plots A and B represent the simulations, with PC1 on the x-axis and PC2 on the y-axis, while a purple-to-yellow color gradient indicates the simulation timeline.

**Figure 10.**
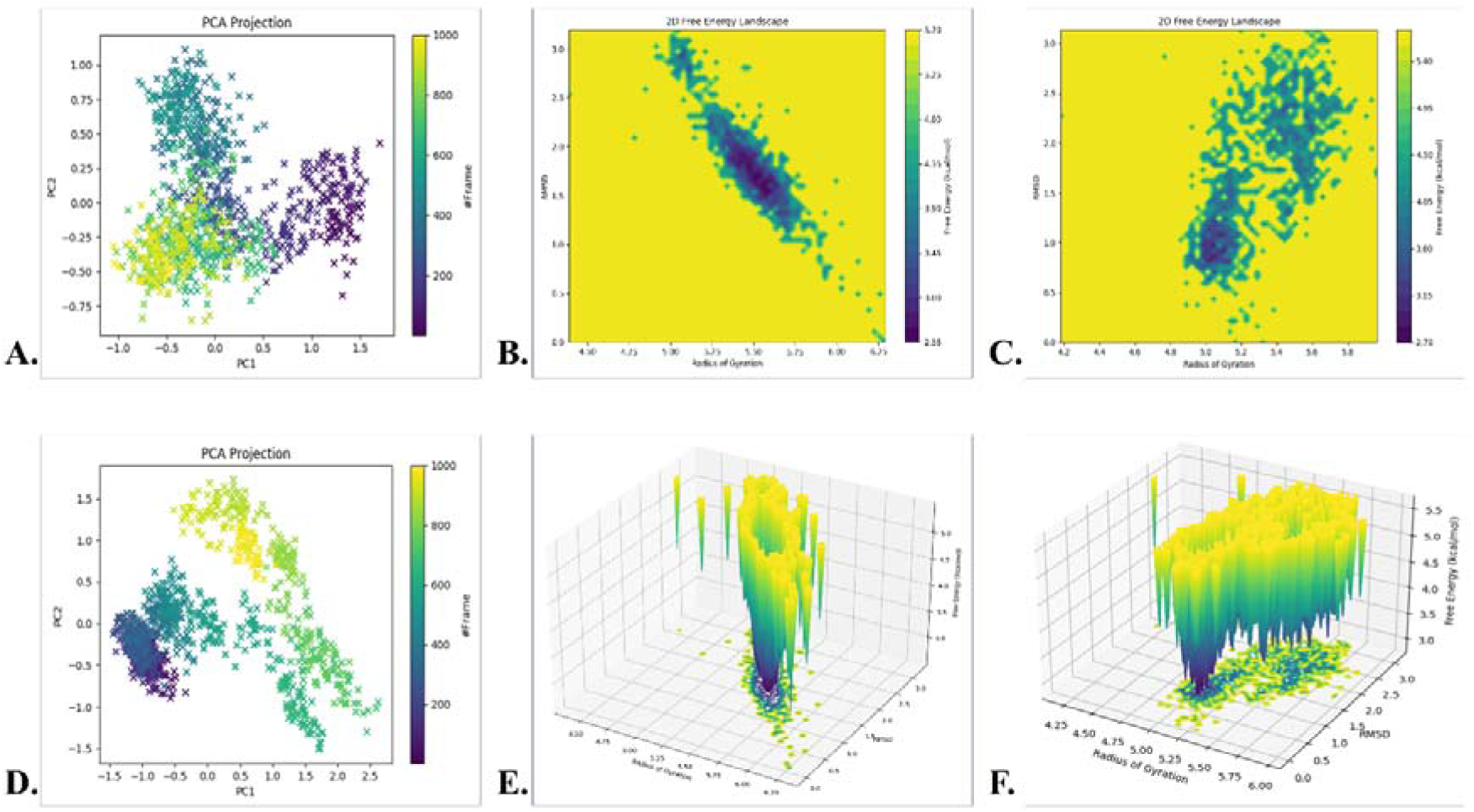
Principal component analysis (PCA) scatter plots, 2D free energy landscape, and 3D free energy surfaces for the molecular dynamics simulations of the complexes MCL1- CNP0161565 (A, B & E) and MCL1- CNP0405001 (D, C & F)

In Plot A, data points cluster densely around the center, indicating constrained conformational sampling and stability of CNP0161565 during the simulation. Most points form a circular cluster near the origin (0,0), suggesting minimal conformational changes and minor flexibility along the positive PC2 axis. The smooth color gradient implies that ligand A does not induce significant conformational changes.

Conversely, Plot D exhibits a broader, more dispersed point distribution, indicating that CNP0405001 allows the MCL1 protein to explore a larger conformational space. This suggests greater flexibility and a diverse range of conformations. Points spread along both PC1 and PC2 axes, with distinct clusters, particularly in the upper right quadrant, indicating different structural arrangements. The less uniform color gradient reflects more abrupt transitions, suggesting that CNP0405001 induces dynamic changes and potentially destabilizes certain conformations.

Comparatively, CNP0161565 keeps the MCL1 protein in a stable conformational state, indicating stronger interactions, while CNP0405001 promotes a more dynamic conformational landscape with multiple metastable states. Thus, CNP0161565 favors stability, while CNP0405001 enhances conformational adaptability, which could impact their biological activity and binding efficiency.

#### 3.7.8. Free Energy Landscape (FEL)

To explore potential conformational shifts in the protein during molecular dynamics simulations, the free energy landscape and Gibbs free energy analysis were conducted. This approach provides insight into how the protein’s structure fluctuates throughout the simulation. Two key variables are employed in this assessment, specifically designed to capture the extent of conformational variation and reveal system-specific characteristics.

The analysis utilized two metrics: the radius of gyration (RoG) and root mean square deviation (RMSD), to map the energy minima for the complexes formed between MCL1 and two different ligands— CNP0161565 and CNP0405001. The study shows a variation in Gibbs free energy (ΔG) ranging from 2.55 to 5.70 kcal/mol for the MCL1- CNP0161565 complex (Figure 9B), and from 2.70 to 5.80 kcal/mol for the MCL1- CNP0405001 complex (Figure 9C). The colour scheme, ranging from yellow to dark purple, indicates the free energy levels, with dark purple areas representing regions of lower energy and higher stability.

The 3D free energy plots (Figures 9E and 9F) further expand on this by providing a more dynamic representation of the energy distribution across different conformational states. These plots reveal a funnel-like structure, especially visible in MCL1- CNP0161565, which indicates a smooth pathway toward a low-energy state. This suggests that the system undergoes minimal conformational changes while maintaining structural stability. MCL1- CNP0405001 displays a more scattered distribution of energy states, indicating a broader range of conformational flexibility. The higher energy values in these regions suggest that the MCL1- CNP0405001 complex may experience greater dynamic movement during the simulation.

## 4. DISCUSSION

Our study advances cancer therapeutics by focusing on MCL1 inhibitors, essential in cellular survival and apoptosis resistance in acute myeloid leukemia (AML). This work employs advanced computational methods, including pharmacophore modelling, QSAR analysis, molecular docking, DFT and molecular dynamics simulations, to identify novel MCL1 inhibitors, paving the way for effective anti-cancer agents.

We utilized pharmacophore modelling to generate models, AAAHHRR_1 and AHHRR, featuring key chemical attributes like hydrogen bond acceptors and aromatic rings. The AAAHHRR_1 model’s high PhaseHypo score indicates its effectiveness in capturing essential binding interactions, crucial for subsequent virtual screening (Jawarkar *et al*., 2022).

The QSAR model, KPLS_Dendritic, demonstrated strong predictive capabilities with high R² and Q² for IC_50_ predictions, although it is essential to address potential limitations by validating predicted values through *in vitro* assays (Yang *et al*., 2023). The dendritic model used in our QSAR analysis captures the complex architecture of molecular structures through its branching framework, which enhances the representation of local and global features. This capability allows for extracting diverse molecular descriptors and topological indices, contributing significantly to understanding how structural variations influence biological activity and improving the predictive power of QSAR models in drug discovery (Teles *et al*., 2022).

Molecular docking identified CNP0161565 and CNP0405001 as top inhibitors with binding affinities of -11.705 kcal/mol and -10.198 kcal/mol, respectively. Key interacting residues include Arg263, Phe270, and Thr266 in CNP0161565 binding. Arg263, mimicking a conserved Asp residue in pro-apoptotic proteins (Mady *et al*., 2018; Marimuthu *et al*., 2020), engages in a cation-π interaction with CNP0161565, emphasizing its role in maintaining the BH3-binding groove’s integrity. Phe270, located in the P2 pocket, is crucial for structural stability (Zhang *et al*., 2023), while Thr266 influences MCL1’s stability and function based on its phosphorylation state (Alhammadi *et al*., 2024; Conage_Pough & Boise, 2018). CNP0405001 interacts through Phe270 and Hie224, demonstrating critical roles in maintaining the hydrophobic binding groove’s conformation (Luna_Vargas & Chipuk, 2016). Phe270 in MCL1’s P2 pocket is crucial for maintaining the hydrophobic binding groove, while Arg263 mimics Asp in pro-apoptotic proteins, emphasizing its role in MCL1’s anti-apoptotic function. Thr266’s phosphorylation status affects MCL1, and Hie224 is vital for conformational integrity.

Agrimonolide is sourced from databases like ZINC-NP (Irwin & Shoichet, 2005) and TCMID (Xue *et al*., 2012), showing potential in breast cancer therapies and a range of biological activities. CNP0405001 (STL544842), derived from InterBioScreen Ltd., is linked to neoflavanoids, exhibiting anti-inflammatory and anticancer properties.

DFT complements our findings by providing accurate predictions of molecular properties and interactions at the quantum mechanical level. It enabled the investigation of electronic structures, optimization of molecular geometries, and evaluation of reaction pathways, essential for understanding ligand-protein interactions and binding affinities (Ye *et al*., 2022).

ADMET analysis highlights solubility, permeability, and metabolic stability challenges, suggesting the need for medicinal chemistry optimization to balance potency and pharmacokinetics. Differences in ADMET profiles between CNP0161565 and CNP0405001 provide insights for structure-activity relationship studies.

MD simulations reveal distinct binding modes: CNP0161565 shows stable RMSD, suggesting rigid binding, while CNP0405001 exhibits higher RMSD, indicating flexibility that may facilitate allosteric interactions. RMSF analysis indicates CNP0161565 induces less protein flexibility, relevant for drug resistance (Garlick & Mapp, 2020). Rg and SASA analyses suggest compact structures can enhance binding but might limit access (Singh *et al*., 2022). PCA results indicate differing conformational preferences, with CNP0161565 stabilizing specific conformations and CNP0405001 allowing more sampling.

Our findings elucidate MCL1 inhibitor structure-activity relationships, positioning CNP0161565 and CNP0405001 as promising candidates for development. A medicinal chemistry program should focus on enhancing solubility, permeability, and metabolic stability while maintaining potency, along with in vivo efficacy studies for biological evaluation. The potential for diverse scaffolds and fragment-based drug discovery approaches offers avenues for improved MCL1 targeting.

This study marks significant progress in MCL1 inhibitor development, integrating pharmacophore modelling, QSAR, DFT, molecular docking, and MD simulations to identify and understand binding mechanisms of promising compounds. Further investigation and optimization are essential for translating these findings into effective clinical therapeutics.

## 5. CONCLUSION

In summary, MCL1 is a critical protein implicated in the progression of various cancers, and targeting it offers a promising therapeutic avenue. The computational framework employed in this study included ligand-based pharmacophore modelling, QSAR analysis, molecular docking, DFT and extensive molecular dynamics simulations. Through this high-throughput virtual screening approach, two lead compounds, CNP0161565 and CNP0405001, were identified as potential natural MCL1 inhibitors. CNP0161565 exhibited superior stability and conformational integrity throughout the 100 ns molecular dynamics simulation, highlighting its potential as an effective therapeutic candidate against MCL1. Additionally, the binding interactions of CNP0161565 with key residues in the MCL1 protein were assessed, revealing favourable interactions that support its role as a selective inhibitor. The calculated binding free energy further substantiates its promise, indicating a robust affinity for the target protein. However, the metabolic pathways, bioavailability, and biotransformation of these compounds post-oral administration remain to be elucidated. Therefore, *in vitro* and *in vivo* studies are recommended to validate the pharmacological activity of CNP0161565 and CNP0405001 against MCL1, establishing their efficacy in a biological context.

## Acknowledgement

We would like to sincerely acknowledge Prof. R Sowdhamini, National Centre for Biological Sciences (NCBS), Tata Institute of Fundamental Research (TIFR), Bangalore, Karnataka, for providing access to the Schrödinger drug discovery suite, which was essential for carrying out this work. I also extend my gratitude to Mr. Varinder Kumar, Assistant Professor of Bioinformatics, Goswami Ganesh Dutta Sanatan Dharma College (GGDSDC), Chandigarh, for granting access to the Gaussian software for Density Functional Theory (DFT) calculations. Additionally, I would like to thank Dr. Abhijit Kayal, Senior Scientist II at Schrödinger, for his invaluable assistance with Free Energy Landscape (FEL) calculations and for his support with the Python scripts used in the analysis.

## Declaration of Competing Interest

The authors report no conflict of interest.

## Ethics Statement

This research did not involve any human participants or animal subjects, and therefore, no ethical approval was required.

## Funding Statement

This research received no external funding.

## Author Contributions

Uddalak Das: Conceptualization, Methodology, Formal Analysis, Writing—Original Draft, Writing—Review & Editing. Tathagata Chanda: Formal Analysis. Jitendra Kumar: Supervision, Project Administration. Anitha Peter: Supervision. All the authors have read and agreed to the published version of the manuscript.

## Notes

### Competing Interest Statement

The authors have declared no competing interest.

