## Supplementary material for "Discovery of Natural MCL1 Inhibitors using Pharmacophore modelling, QSAR, Docking, ADMET, Molecular Dynamics, and DFT Analysis": Supplimentary Information

**SUPPLEMENTARY DATA**

**Table 1** Pharmacophore model evaluation using multiple scoring metrics for all the 40 generated pharmacophore models, sorted based on PhaseHypo Score

| **Sl. No.** | **Title** | **Survival Score** | **Site Score** | **Vector Score** | **Volume Score** | **Selectivity Score** | **Num Matched** | **Inactive Score** | **Adjusted Score** | **BEDROC Score** | **PhaseHypo Score** |
| --- | --- | --- | --- | --- | --- | --- | --- | --- | --- | --- | --- |
| 1 | **AAAHHRR_1** | **8.263797** | **0.719794** | **0.928613** | **0.638179** | **3.431904** | **351** | **0** | **8.263797** | **0.999919** | **1.495728** |
| 2 | AAHHHRR_1 | 8.197675 | 0.662387 | 0.932748 | 0.674602 | 3.376488 | 356 | 0 | 8.197675 | 0.999924 | 1.491762 |
| 3 | AAHHHRR_2 | 8.197528 | 0.699191 | 0.933044 | 0.68935 | 3.335614 | 347 | 0 | 8.197528 | 0.999892 | 1.49175 |
| 4 | AAHHHRR_3 | 8.095258 | 0.634369 | 0.921315 | 0.670086 | 3.320484 | 354 | 0 | 8.095258 | 0.999892 | 1.485618 |
| 5 | AAHHHRR_4 | 8.058613 | 0.575728 | 0.92433 | 0.631686 | 3.381563 | 351 | 0 | 8.058613 | 0.998918 | 1.482416 |
| 6 | AAHHHRR_5 | 8.023264 | 0.627559 | 0.941536 | 0.622824 | 3.310206 | 332 | 0 | 8.023264 | 0.999907 | 1.481298 |
| 7 | AAHHHRR_6 | 7.901934 | 0.590948 | 0.91881 | 0.566041 | 3.290841 | 343 | 0 | 7.901934 | 0.999022 | 1.473114 |
| 8 | AAHHHRR_7 | 7.870857 | 0.554698 | 0.90199 | 0.524595 | 3.339345 | 355 | 0 | 7.870857 | 0.999924 | 1.472154 |
| 9 | AAHHHRR_8 | 7.856944 | 0.55616 | 0.907043 | 0.521484 | 3.322028 | 355 | 0 | 7.856944 | 0.999923 | 1.471314 |
| 10 | AAHHHRR_9 | 7.839064 | 0.521655 | 0.864856 | 0.501031 | 3.401294 | 355 | 0 | 7.839064 | 0.999901 | 1.470246 |
| 11 | AAHHRR_1 | 7.721477 | 0.753884 | 0.917741 | 0.652305 | 2.852241 | 351 | 0 | 7.721477 | 0.999899 | 1.46319 |
| 12 | AAHHRR_2 | 7.624613 | 0.73101 | 0.957591 | 0.635628 | 2.751381 | 354 | 0 | 7.624613 | 0.999925 | 1.457376 |
| 13 | AHHHRR_1 | 7.612095 | 0.741835 | 0.908344 | 0.692281 | 2.728056 | 348 | 0 | 7.612095 | 0.999903 | 1.456626 |
| 14 | AHHHRR_2 | 7.596248 | 0.703748 | 0.909681 | 0.673774 | 2.760043 | 354 | 0 | 7.596248 | 0.999897 | 1.455672 |
| 15 | AAHHRR_3 | 7.559254 | 0.768061 | 0.93987 | 0.671633 | 2.624595 | 359 | 0 | 7.559254 | 0.999918 | 1.453458 |
| 16 | AHHHRR_3 | 7.541991 | 0.706095 | 0.890478 | 0.675034 | 2.720155 | 355 | 0 | 7.541991 | 0.999697 | 1.45222 |
| 17 | AAAHHR_1 | 7.52784 | 0.707569 | 0.927743 | 0.636367 | 2.710854 | 351 | 0 | 7.52784 | 0.999732 | 1.451368 |
| 18 | AHHHRR_4 | 7.494322 | 0.7197 | 0.962694 | 0.686135 | 2.584213 | 348 | 0 | 7.494322 | 0.999963 | 1.449658 |
| 19 | AHHHRR_6 | 7.457768 | 0.693662 | 0.957181 | 0.659389 | 2.592441 | 359 | 0 | 7.457768 | 0.999923 | 1.447368 |
| 20 | AHHHRR_5 | 7.469146 | 0.606999 | 0.900188 | 0.641984 | 2.773432 | 352 | 0 | 7.469146 | 0.998813 | 1.446946 |
| 21 | AHHRR_1 | 7.150725 | 0.779231 | 0.941028 | 0.646059 | 2.234178 | 355 | 0 | 7.150725 | 0.99998 | 1.429042 |
| 22 | AHHRR_2 | 7.108378 | 0.828026 | 0.929803 | 0.663592 | 2.130654 | 360 | 0 | 7.108378 | 0.999928 | 1.426404 |
| 23 | AAHHR_1 | 7.099827 | 0.743795 | 0.944888 | 0.644811 | 2.216105 | 355 | 0 | 7.099827 | 0.998735 | 1.424688 |
| 24 | AAHHR_2 | 6.988201 | 0.72601 | 0.961591 | 0.631215 | 2.119156 | 355 | 0 | 6.988201 | 0.999951 | 1.419292 |
| 25 | AHHRR_3 | 6.988987 | 0.755034 | 0.892667 | 0.663756 | 2.121228 | 360 | 0 | 6.988987 | 0.999645 | 1.41894 |
| 26 | HHHRR_1 | 6.976989 | 0.759443 | 0.936535 | 0.682668 | 2.048115 | 355 | 0 | 6.976989 | 0.999905 | 1.41852 |
| 27 | AHHRR_4 | 6.974462 | 0.798922 | 0.96475 | 0.677682 | 1.976806 | 360 | 0 | 6.974462 | 0.999962 | 1.41847 |
| 28 | AHHHR_1 | 6.95508 | 0.761456 | 0.965554 | 0.673866 | 2.002754 | 356 | 0 | 6.95508 | 0.99996 | 1.417306 |
| 29 | AAHHR_3 | 6.934755 | 0.76641 | 0.939018 | 0.66771 | 2.005314 | 360 | 0 | 6.934755 | 0.999965 | 1.416088 |
| 30 | AHHHR_2 | 6.931828 | 0.708688 | 0.962495 | 0.662709 | 2.046485 | 356 | 0 | 6.931828 | 0.999968 | 1.415908 |
| 31 | AHHR_2 | 6.650105 | 0.862415 | 0.975796 | 0.659705 | 1.595886 | 360 | 0 | 6.650105 | 0.999995 | 1.399006 |
| 32 | AHHR_1 | 6.652005 | 0.819338 | 0.955081 | 0.634417 | 1.686867 | 360 | 0 | 6.652005 | 0.999728 | 1.39882 |
| 33 | HHRR_1 | 6.594106 | 0.869046 | 0.951548 | 0.670192 | 1.547018 | 360 | 0 | 6.594106 | 0.999958 | 1.395646 |
| 34 | HHRR_2 | 6.501209 | 0.788951 | 0.925992 | 0.688308 | 1.541655 | 360 | 0 | 6.501209 | 0.999851 | 1.389972 |
| 35 | HHRR_3 | 6.487773 | 0.877334 | 0.91522 | 0.591034 | 1.547883 | 360 | 0 | 6.487773 | 0.999766 | 1.389068 |
| 36 | AHRR_1 | 6.484614 | 0.84611 | 0.930475 | 0.6638 | 1.487927 | 360 | 0 | 6.484614 | 0.999443 | 1.388476 |
| 37 | HHHR_1 | 6.474037 | 0.876538 | 0.990225 | 0.643186 | 1.407785 | 360 | 0 | 6.474037 | 0.999965 | 1.38844 |
| 38 | AHHR_3 | 6.470265 | 0.762609 | 0.925094 | 0.644951 | 1.587383 | 355 | 0 | 6.470265 | 0.999943 | 1.388118 |
| 39 | AHRR_3 | 6.468986 | 0.781844 | 0.941123 | 0.630286 | 1.559431 | 360 | 0 | 6.468986 | 0.999762 | 1.38794 |
| 40 | AHRR_2 | 6.48351 | 0.853788 | 0.93106 | 0.647673 | 1.494686 | 360 | 0 | 6.48351 | 0.996701 | 1.38571 |

**Table 2** Performance metrics of 10 QSAR models**.**

| Sl. No. | Model Code | Score | S.D. | R^2 | RMSE | Q^2 | Q^2 MW (Null Hypothesis) |
| --- | --- | --- | --- | --- | --- | --- | --- |
| 1 | **kpls_dendritic_1** | **0.8997** | **0.0022** | **0.8997** | **0.0021** | **0.8977** | **-0.0108** |
| 2 | kpls_dendritic_2 | 0.8942 | 0.0022 | 0.8941 | 0.0022 | 0.8925 | 0.0268 |
| 3 | kpls_dendritic_30 | 0.8932 | 0.0022 | 0.8917 | 0.0020 | 0.9077 | 0.0255 |
| 4 | kpls_linear_30 | 0.8866 | 0.0022 | 0.8972 | 0.0022 | 0.8907 | 0.0255 |
| 5 | kpls_dendritic_40 | 0.8773 | 0.0022 | 0.8923 | 0.0023 | 0.8860 | 0.0148 |
| 6 | kpls_linear_40 | 0.8560 | 0.0022 | 0.8998 | 0.0024 | 0.8781 | 0.0148 |
| 7 | kpls_linear_2 | 0.8543 | 0.0022 | 0.8924 | 0.0024 | 0.8690 | 0.0268 |
| 8 | kpls_dendritic_34 | 0.8103 | 0.0022 | 0.8940 | 0.0026 | 0.8460 | 0.0272 |
| 9 | kpls_radial_13 | 0.8089 | 0.0022 | 0.8914 | 0.0026 | 0.8417 | 0.0348 |
| 10 | kpls_radial_35 | 0.8028 | 0.0023 | 0.8823 | 0.0027 | 0.8422 | 0.0026 |

**Table 3** Physiochemical properties of the two screened compounds

| Properties | CNP0161565 | CNP0405001 |
| --- | --- | --- |
| Molecular Weight | 476.47 | 454.50 |
| Dipole Moment | 2.298 | 7.354 |
| Density | 1.038 | 0.983 |
| H-bond donors | 5 | 1 |
| H-bond acceptors | 10 | 7 |
| Number of rotatable bonds | 7 | 7 |
| Number of rings | 4 | 5 |
| Number of atoms in the biggest ring | 10 | 10 |
| Number of heteroatoms | 10 | 7 |
| Formal Charge | 0 | 0 |
| Number of rigid bonds | 24 | 29 |
| Sterio Centres | 6 | 5 |
| SASA | 735.722 | 769.281 |
| FOSA | 320.367 | 400.191 |
| FISA | 242.583 | 141.247 |
| PISA | 172.772 | 227.842 |
| WPSA | 0 | 0 |
| Total solvent accessible volume (in Å^3^) | 1366.838 | 1422.751 |
| Polarizability (in Å^3^) | 42.492 | 48.466 |
| Hexadecane/Gas partition coefficient | 14.766 | 14.096 |
| Octanol/Gas partition coefficient | 26.996 | 20.074 |
| Water/Gas partition coefficient | 20.085 | 8.939 |
| Octanol/Water partition coefficient | 0.864 | 4.701 |
| Number of non-conjugated amine groups | 0 | 0 |
| Number of amidine and guanidine groups | 0 | 0 |
| Number of carboxylic acid groups | 0 | 0 |
| Number of non-conjugated amide groups | 0 | 0 |
| Number of reactive functional groups | 2 | 1 |
| PM3 calculated ionization potential (negative of HOMO energy) (in eV) | 8.876 | 8.779 |
| PM3 calculated electron affinity (negative of LUMO energy) (in eV) | 0.698 | 1.04 |

**Figure 4** Medicinal Chemistry properties of the two screened compounds

| Properties | CNP0161565 | CNP0405001 |
| --- | --- | --- |
| Synthetic Accessibility Score | Easy | Easy |
| MCE-18 Value | 85.0 | 90.222 |
| PAINS | 0 alert | 0 alert |
| Brenk | 0 alert | 1 alert |
| ALARM NMR Rule | 2 alert | 2 alert |
| BMS Rule | 0 alert | 0 alert |
| Chelator Rule | 0 alert | 0 alert |
| Lipinski’s Rule of 5 violations | 0 violations | 0 violations |
| Golden Triangle | Accepted | Accepted |
| Pfizer Rule | Accepted | Accepted |

**Figure 5** ADMET properties of the two screened compounds

| **Properties** | | **CNP0161565** | **CNP0405001** |
| --- | --- | --- | --- |
| **A** | Water solubility (log mol/L) | -3.689 | -5.098 |
|  | Caco-2 permeability (Papp in 10^-6^ cm/s) | -0.131 | 1.032 |
|  | Human intestinal absorption (in %) | 40.636 | 99.699 |
|  | Skin Permeability (log K_p_) | -2.735 | -2.735 |
|  | Human oral absorption (in %) | 62.353 | 100 |
|  | P-glycoprotein substrate | Yes | No |
|  | P-glycoprotein I inhibitor | Yes | Yes |
|  | P-glycoprotein II inhibitor | No | Yes |
|  | MDCK Permeability (in nm/sec) | 0.00 | 0.00 |
| **D** | Volume of distribution at steady state (in log L/kg) | 0.766 | -0.905 |
|  | Plasma protein binding (in %) | 81.2 | 98.8 |
|  | Fraction of unbound plasms | 18.7 | 0.9% |
|  | BBB Permeability (in logBB) | -1.399 | -0.678 |
|  | CNS permeability (in logPS) | -4.184 | -3.11 |
| **M** | CYP2D6 substrate | No | No |
|  | CYP3A4 substrate | No | Yes |
|  | CYPIA2 inhibitor | No | No |
|  | CYP2C19 inhibitor | No | Yes |
|  | CYP2C9 inhibitor | No | Yes |
|  | CYP2D6 inhibitor | No | No |
|  | CYP3A4 inhibitor | No | Yes |
| **E** | Total Clearance (in log ml/min/kg) | 0.729 | 0.35 |
|  | Renal OCT2 substrate | No | No |
| **T** | AMES toxicity | Yes | No |
|  | Max. tolerated dose (human) | 0.315 | 0.226 |
|  | hERG I inhibitor | No | No |
|  | hERG II inhibitor | No | Yes |
|  | Oral Rat Acute Toxicity (LD50) (mol/kg) | 3.461 | 2.935 |
|  | Oral Rat Chronic Toxicity (LOAEL) (log mg/kg/day) | 4.385 | 1.429 |
|  | Skin Sensitisation | No | No |
|  | Hepatotoxicity | No | No |
|  | *T Pyriformis* toxicity (µg/L) | 0.285 | 0.286 |
|  | Minnow toxicity (log mM) | 2.751 | -2.197 |
|  | FDA Maximum Recommended Daily Dose | 0.302 | 0.731 |
|  | Carcinogenicity | No | No |
|  | Eye corrosion | No | No |
|  | Eye irritation | No | No |
|  | Respiratory toxicity | No | No |
|  | Predicted GHS Toxicity Class | 6 | 4 |

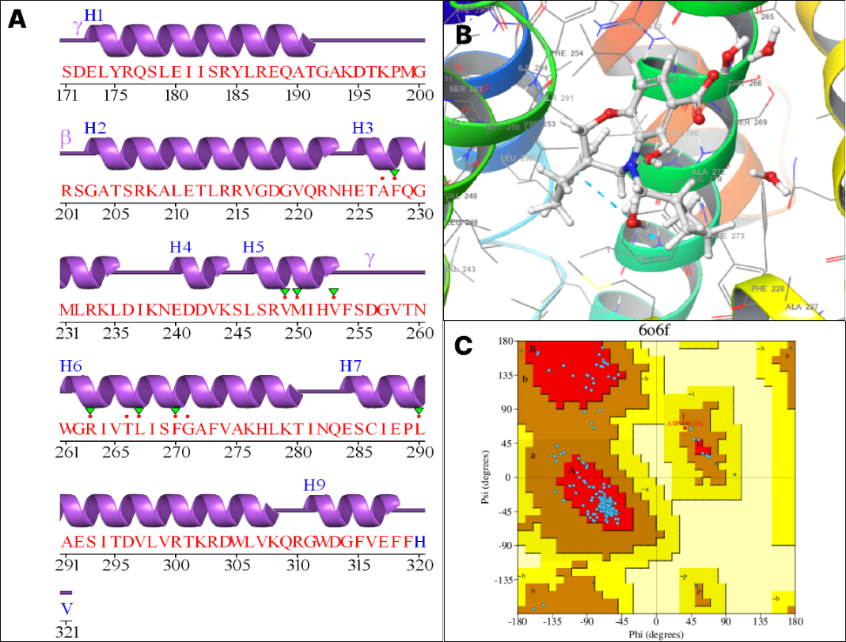

**Figure 1** The secondary structure of the amino acid sequence of protein 6O6F(A) (generated by PDBsum); B: The 3D structure of the MCL1 (Chain A) protein; C: Ramachandran plot of the protein. The colour red indicates Iow-energy regions, yellow - allowed regions, pale yellow - generously allowed regions, and white - disallowed regions.

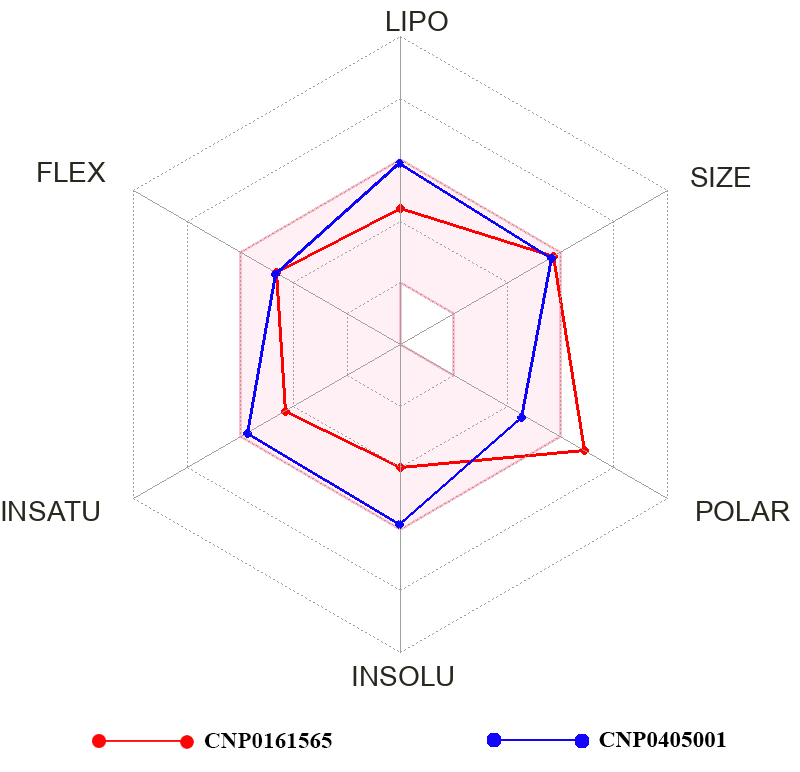

**Figure 2** A radar plot (generated from SwissADME) comparing the lipophilicity (LIPO), flexibility (FLEX), saturation (INSATU), polarity (POLAR), and insolubility (INSOLU) properties of CNP0161565 and CNP0405001. The faded blue region marks the optimal drug parameters.

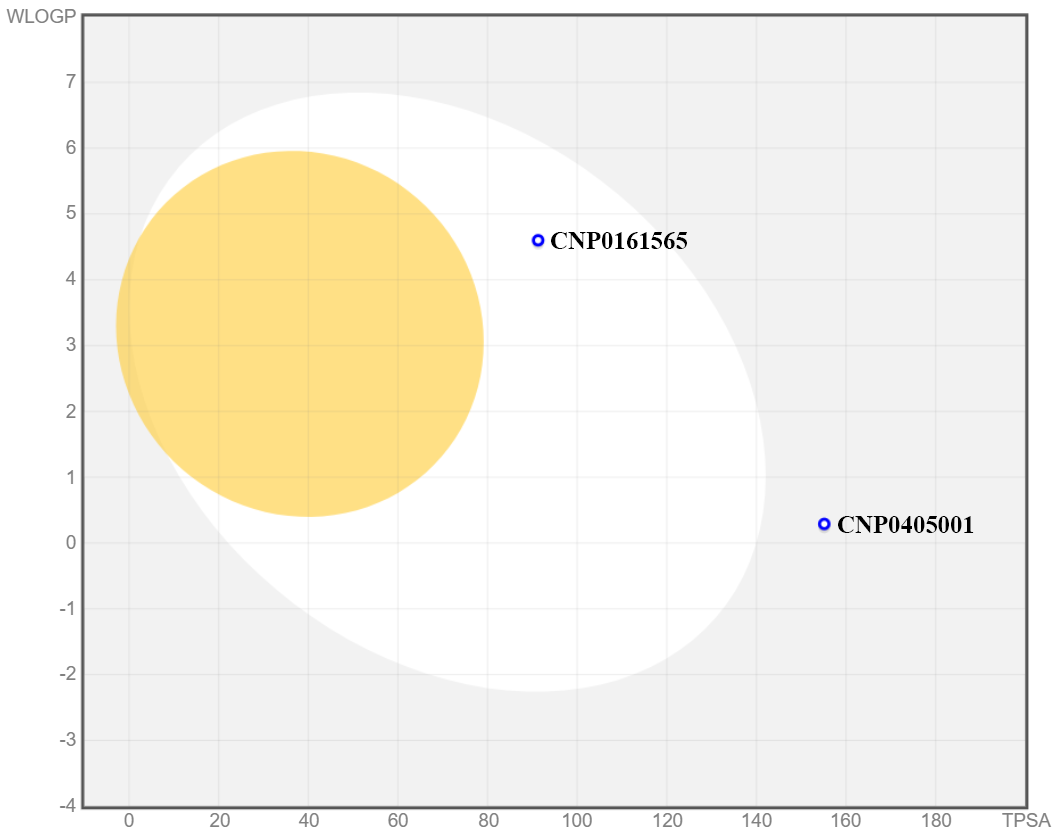

**Figure 3** A BOILED-Egg plot (generated grom SwissADME) illustrating the predicted gastrointestinal absorption and brain penetration of two compounds, CNP0161565 and CNP0405001, based on their WLOGP and TPSA values
